# Role of Uridine Phosphorylase 1 (UPP1) in Pancreatic β-Cell Physiology and Its Dysregulation in Obesity and Type 2 Diabetes

**DOI:** 10.64898/2026.09.24.754194

**Authors:** Yunqian Peng, Lu Ding, Qiutang Xiong, Ling Fu, Chelsea Jang, Yin Wang, Ignacio Norambuena-Soto, Kevin Wang, Gemin Lu, Erika M. McCown, Eunjin Oh, Lin Tan, Jeffrey S. Isenberg, Ping Wang, Philipp E. Scherer, Zhao V. Wang, Adolfo Garcia-Ocana, Debbie C. Thurmond, Yingfeng Deng

## Abstract

**Aims/hypothesis:** Fasting plasma uridine concentrations are elevated in obese mice and in individuals with type 2 diabetes (T2D), suggesting that uridine may function as a metabolic signal that adjusts insulin secretion to changes in metabolic demand. Here, we tested this hypothesis by determining whether uridine potentiates glucose-stimulated insulin secretion (GSIS) in mouse and human islets and by identifying uridine phosphorylase 1 (UPP1), the principal enzyme responsible for systemic uridine catabolism, as a key mediator of this effect.

**Methods:** Whole-body *Upp1* knockout (KO) and β-cell-specific *Upp1* knockout (KO^β^) mice were used to determine the requirement for UPP1 in uridine-potentiated glucose-stimulated insulin secretion (GSIS) and glucose-stimulated MAPK activation. Leptin-deficient and diet-induced obese mouse models were used to assess the impact of obesity on uridine responsiveness. UPP1 protein abundance and subcellular localization were examined in islets from obese mice and human donors spanning a range of glycemic control. Integrated single-nucleus RNA sequencing and ATAC sequencing were performed on wild-type and *Upp1* KO mouse islets to characterize β-cell molecular remodeling associated with UPP1 deficiency and altered responses to uridine.

**Results:** Uridine potentiated GSIS in islets from chow-fed and high-fat diet-fed mice, but not in leptin-deficient obese mice. Uridine also potentiated GSIS in human islets, although this effect progressively declined with worsening glycemic control. In mouse islets, uridine potentiation was accompanied by transient activation of the MAPK pathway. Both uridine-potentiated GSIS and uridine-enhanced glucose-stimulated MAPK activation were abolished in islets from *Upp1* KO and KO^β^ mice. The requirement for a transient MAPK signaling was further supported by pharmacological inhibition of MEK and constitutive activation of MEK1. Leptin-deficient islets exhibited impaired glucose-induced MAPK activation together with increased UPP1 protein expression. In human islets, UPP1 protein abundance positively correlated with donor HbA_1c_ and displayed heterogeneous subcellular localization. Integrated single-nucleus multiomic analysis identified UPP1 as an important regulator of β-cell identity, function, and transcriptional responses to uridine.

**Conclusions:** These findings establish UPP1 as a key regulator linking pyrimidine metabolism to β-cell function and identify impaired uridine responsiveness as a potential contributor to and/or consequence of β-cell dysfunction during the progression of T2D.

**Research in Context:** What is already known about this subject?

- Circulating uridine concentrations are elevated in obesity and type 2 diabetes (T2D)
- Uridine phosphorylase 1 (UPP1) is the principal enzyme responsible for uridine catabolism and plays a key role in regulating systemic uridine homeostasis.

What is the key question?

- Does uridine regulate pancreatic β-cell function, what role does UPP1 play in mediating this effect, and how is this pathway altered during obesity and the progression of T2D?

What are the new findings?

- Uridine potentiates glucose-stimulated insulin secretion (GSIS) in a UPP1-dependent manner, accompanied by transient activation of the MAPK signaling pathway.
- Uridine potentiation of GSIS is preserved in diet-induced obesity but abolished in leptin-deficient obesity, and progressively declines in human islets with worsening glycemic control.
- Disrupted MAPK pathway and aberrant increased UPP1 protein expression are associated with impaired uridine potentiation of GSIS in obesity and T2D, whereas genetic deletion of *Upp1* affects β-cell identity, β-cell function, and the transcriptional responses to uridine.

How might this impact clinical practice in the foreseeable future?

- These findings identify UPP1 as a previously unrecognized regulator of β-cell function that links pyrimidine metabolism to insulin secretion. Targeting the UPP1–uridine pathway may represent a new strategy to preserve β-cell function or improve insulin secretory capacity during the progression of T2D.

## Introduction

Uridine is the most abundant circulating nucleoside and participates in a wide range of physiological processes [1, 2]. We previously demonstrated that circulating uridine concentrations are regulated by nutritional status, increasing during fasting in both mice and humans [3]. This fasting–feeding regulation has subsequently been confirmed in healthy individuals [4]. In contrast, individuals with severe obesity exhibit elevated postprandial rather than reduced circulating uridine concentrations [5], suggesting that obesity disrupts systemic uridine homeostasis. However, whether excess uridine contributes to the development of obesity-associated metabolic dysfunction and diabetes remains unclear.

Elevated plasma uridine concentrations were first reported in individuals with hypertension and have since been associated with metabolic disease [6]. Fasting plasma uridine positively correlates with fasting insulin levels and the homeostasis model assessment of insulin resistance [7] and independently predicts the risk of T2D [8]. Mechanistically, excessive uridine impairs insulin signaling through enhanced protein O-GlcNAcylation [9–12], and chronic dietary uridine supplementation induces hepatic steatosis and insulin resistance in mice [13], supporting a causal role for elevated uridine in metabolic dysfunction.

Uridine phosphorylase 1 (UPP1), the first and rate-limiting enzyme in uridine catabolism [14], is a major regulator of systemic uridine homeostasis. Whole-body deletion of *Upp1* increases circulating uridine concentrations by more than sixfold [15], whereas hepatocyte-specific overexpression of *Upp1* markedly lowers circulating uridine levels [16]. These findings identify UPP1 as a key determinant of circulating uridine; however, whether impaired UPP1 activity contributes to the elevation of plasma uridine during obesity and T2D remains unknown. Our recent analysis of biospecimens from the Diabetes Prevention Trial-Type 1 showed that alterations in fasting uridine concentrations preceded the onset of type 1 diabetes (T1D) and that fasting uridine concentrations at the screening visit had predictive value for T1D development [17], raising the possibility that uridine participates in the regulation of β-cell function. In this study, we investigated how uridine regulates β-cell function through UPP1 and how this pathway is altered during obesity and T2D progression.

## Methods

### Mouse experiments

Adult C57BL/6N wild-type (WT) mice and other mouse strains were housed in individually ventilated cages under controlled temperature on a 12-h light/12-h dark cycle with ad libitum access to food and water. Unless otherwise indicated, mice were maintained on a standard rodent chow diet (2916, Teklad) and switched to experimental diets at 10-15 weeks of age. A high-fat diet (HFD; D12492, Research Diet) was used to induce obesity. Doxycycline-containing chow (600 mg/kg, Bio-serv) was used to induce β-cell-specific deletion of *Upp1* through the Tet-On system. Whole-body *Upp1* knockout (KO) mice were rederived from frozen sperm obtained from the NIH-supported Mutant Mouse Resource & Research Centers, which had been originally deposited by Giuseppe Pizzorno [15]. Leptin signaling-deficient *ob/ob* mice and tet-responsive element-driven Cre (TRE-Cre) mice were obtained from the Jackson Laboratory and have been described previously [3, 18]. Mouse insulin promoter driven-rtTA (MIP-rtTA) mice were generated in the Scherer lab as previously described [19]. *Upp1*^f/f^ mice were generated from an embryonic stem cell clone obtained from the Knock Out Mouse Project (University of California, Davis). All mouse lines were backcrossed onto the C57BL/6N background for more than nine generations before being intercrossed to generate inducible β-cell-specific *Upp1* KO (*Upp1* KO^β^) mice. Sex– and age-matched littermates homozygous for *Upp1*^f/f^ but lacking either MIP-rtTA, TRE-Cre, or both transgenes served control mice. Mice were killed for islet isolation at the indicated ages. Oral glucose tolerance tests (OGTTs) were performed in mice after a 4–6 h fast, as previously described [20]. Glucose (Sigma) was administrated by oral gavage as a 25% solution (w/vol) at 2.5 g/kg bodyweight. Blood samples were collected by tail nick into heparinized capillary tubes (Fisher Scientific) at 0, 15, 30, 60, and 120 min. Blood glucose concentrations were measured at each time point using a glucometer (Ascensia Contour next). Plasma insulin concentrations were measured using a mouse insulin ELISA kit (Crystal Chem). All animal procedures were approved by the Institutional Animal Care and Use Committees of the City of Hope and the University of Texas Southwestern Medical Center.

### Islet functional studies

Mouse islet were isolated by collagenase digestion of the pancreas, as previously described [21]. Islets were cultured in RPMI 1640 medium (Fisher Scientific) supplemented with 10% fetal bovine serum (FBS), 100 U/ml penicillin, 0.1 mg/ml streptomycin, and 8 mM glucose. Static glucose-stimulated insulin secretion (GSIS) assays were performed as previously described [22]. Briefly, 10 islets (150–200 μm in diameter) were transferred to 12-well plates containing 1 ml RPMI 1640 supplemented with 3 mM glucose, 10% FBS, 100 U/ml penicillin, and 0.1 mg/ml streptomycin and preincubated for 2 h. The islets were then incubated for 1 h in Krebs-Ringer bicarbonate HEPES (KRBH) buffer containing 3 mM glucose, followed by incubation for an additional 1 h in KRBH buffer containing 16.7 mM glucose to measure insulin secretion.

Human islets were obtained from the Integrated Islet Distribution Program and the City of Hope Southern California Islet Cell Resource Center. Upon receipt, islets designated for GSIS studies were cultured overnight in RPMI 1640 medium supplemented with 10% FBS and 5.6 mM glucose before being subjected to static GSIS assays as previously described [23].

Glucose, uridine, dilazep dihydrochloride (Bio-techne), and PD0325901 (Selleckchem) were added at the indicated concentrations as previously described [24]. Freshly isolated mouse islets were incubated with adenovirus expressing GFP (Ad-GFP) or constitutively active MEK1 (Ad-CaMEK) in RPMI 1640 medium containing 8 mM glucose for 48 h, as previously described [25, 26].

Dynamic GSIS assays were performed using an islet perifusion system (Biorep Technologies), as previously described [27, 28]. For each assay, 50 islet equivalents were handpicked and loaded into perifusion chambers. Islets were equilibrated for 90 min with baseline HEPES-buffer solution and then subjected to the following perifusion sequence: 2.8 mM glucose for 10 min, 16.7 mM glucose for 35 min, 3 mM glucose for 20min, 30 mM KCl containing 2.8 mM glucose for 5 min, and 2.8 mM glucose for10 min. Perifusate samples were collected at 1-min intervals. At the end of the assay, islets were collected and lysed in 500 μl of 1 M acetic acid containing protease inhibitors to measure total insulin content. Insulin secretion was normalised to total islet insulin content.

Mitochondrial oxygen consumption rate (OCR) was measured using a Seahorse XFe24 Analyzer (Agilent Technologies), as previously described [29]. For each assay, 50 islet equivalent were handpicked and loaded into each well of an XF24 Islet Capture Microplate. islets were maintained in Seahorse XF assay medium containing 3 mM glucose and 1% FBS and equilibrated for 1 h at 37°C in a non-CO₂ incubator. Five OCR measurements were obtained under each condition. After baseline measurement at 3 mM glucose, the following compounds were sequentially injected: glucose to a final concentration of 16.8 mM, 5 µM oligomycin, 1 μM FCCP [carbonyl cyanide-4-(trifluoromethoxy)phenylhydrazone], and 5 μM rotenone plus 5 μM antimycin A. Raw data were analyzed using Wave software (version 2.6; Agilent Technologies), and OCR was normalised to islet equivalent.

For calcium response analysis, WT mouse islets were infected with 1 × 10⁸ PFU of adenovirus expressing CMV-GCaMP5G for 12 h, as previously described [30]. The medium was then replaced with fresh complete medium, and islets were cultured for an additional 36 h before overnight treatment with or without uridine. For Ca^2+^ imaging, islets were transferred to KRBH containing 3 mM glucose and preincubated for 1.5 h. Islets were then allowed to attach for 0.5 h to coverslips coated with poly-D-lysine and laminin (Cell Applications) and imaged at 37°C in an in-house recording chamber using a ZEISS LSM900 confocal microscope. Image sequences were acquired at 30ms-interval and analyzed using ZEISS LSM900 image analysis software for cell identification, background subtraction, and quantification of fluorescence intensity within defined regions of interest (ROIs). Background-subtracted fluorescence intensity (*F*) was normalized to baseline fluorescence (*F₀*) and calcium responses were expressed as *F/F_0_*. The mean *F/F_0_* between 30 and 90 s was calculated for each cell and used to compare response between uridine-treated and untreated islets.

For uridine uptake assay, islets isolated from WT and *ob/ob* mice were cultured overnight in complete RPMI 1640 medium containing 8 mM glucose. Islets were then incubated for 1 h under GSIS condition in KRBH buffer containing 16.7 mM glucose and 100 μM [U-^15^N_2_, U-^13^C_ti_]uridine (Cambridge Isotope Laboratories). Each *ob/ob* sample comprised 250 islets isolated from one mouse, whereas each WT sample comprised 500 islets from 4 mice. Different numbers of WT and *ob/ob* islets were used to balance the total islet biomass because *ob/ob* islets were larger than WT islets. Following incubation, all islets and the corresponding culture medium were collected for metabolomic analysis. An equal volume of blank medium was analyzed to verify the purity and starting abundance of [U-^15^N_2_, U-^13^C_9_]uridine. Metabolomic analysis was performed by the MD Anderson Cancer Center Metabolomics Core. Islet and medium extracts were analyzed by liquid chromatography-ultra-high-resolution mass spectrometry using a Thermo Orbitrap Exploris 240 Mass Spectrometer operated in positive/negative electrospray-ionization polarity-switching mode at a resolution of 120,000. Raw data files were processed using Skyline-Daily, and the abundance of each isotopologue was calculated using AccuCor2, as previously described [31].

### Islet single-nucleus multiomic sample preparation and analysis

Mouse islets were processed for paired single-nucleus RNA sequencing (snRNA-seq) and single-nucleus assay for transposase-accessible chromatin sequencing (snATAC-seq), as previously described [32]. Briefly, islets were dispersed into single-cell suspensions, and sample viability was confirmed to exceed 90% to minimize the carryover of dead cells. Nuclei were isolated from the single-cell suspensions according to the 10x Genomics Multiome ATAC + Gene Expression protocol (CGOOO338). Samples were collected, washed with PBS containing 0.04% BSA, lysed for 4 min in pre-chilled lysis buffer, washed three times with wash buffer, and resuspended in 10x Genomics nuclei buffer at a concentration of 3,000–5,000 nuclei/μl. Nuclei were processed using the 10x Genomics Chromium platform, with a target recovery of 7,000– 1,0000 nuclei per sample. Single-nucleus RNA and ATAC libraries were prepared using the Chromium Single Cell Multiome ATAC + Gene Expression v1 kit according to the manufacturer’s instructions. The quality of all libraries was assessed using an Agilent TapeStation system. RNA and ATAC libraries were sequenced using an Illumina NovaSeq 6000 system.

Multiomic sequencing files (FASTQ) were processed and analyzed using Cell Ranger ARC v2.0. Sequencing reads were aligned to the mouse reference genome (mm10), and gene expression was quantified using the corresponding gene annotation. The resulting datasets were analyzed using Seurat v4.4.0 and Signac v1.11.0 in R v4.4.2. For the snATAC-seq data, peaks were called using MACS2 and annotated based on the mm10 reference genome using EnsDb.Mmusculus.v79 (v2.99.0). Low-quality nuclei, nuclei with insufficient sequencing depth, and putative multiplets were excluded based on RNA and ATAC fragment counts and mitochondrial transcript abundance. Specifically, nuclei were retained if they had 3,000–25,000 RNA counts and 2,000–70,000 ATAC counts, with mitochondrial transcripts representing ≤25% of the total RNA counts. Details for sample preparation and data analysis was provided in ESM method.

### Quantitative PCR, Immunoblotting, and Immunofluorescence

Total RNA was extracted from whole islets and the relative mRNA level of each target gene was calculated using the 2(−△△Ct) method. Primer sequences are provided in ESM Fig. 2.

For immunoblotting, islets were lysed in RIPA buffer supplemented with protease and phosphatase inhibitors (Thermo Scientific). Blots were quantified by Odyssey image analysis software. Antibodies were listed in ESM method.

For immunofluorescence, mouse pancreata were fixed in 4% formalin and embedded in paraffin. Paraffin processing, embedding, and sectioning of human and mouse pancreata were performed by the AR-DMRI Laboratory & Translational Service Centers and the Pathology Core, respectively, at City of Hope. Antibodies were listed in ESM method.

### Studies involving human cadaveric tissue

Samples originate from de-identified cadaveric donors and are institutional review board exempt.

### Statistical analyses

Statistical analyses were performed using GraphPad Prism Software 10.3.0. The statistical tests used for each analysis are specified in the corresponding figure legends. Data are presented as mean ± SEM. A *p*-value <0.05 was considered statistically significant.

## Results

### Uridine potentiates glucose-stimulated insulin secretion in mouse and human islets

Previous studies have used 100 µM uridine to investigate lipid metabolism in primary hepatocytes [16], while uridine concentrations of 200 μM [33] and even 1 mM [34] maintained for 24 h have not been reported to cause detectable cytotoxicity in cultured cells [34]. RPMI 1640 medium contains no uridine, whereas supplementation with 10% fetal bovine serum provides 2–3 µM uridine [18]. Therefore, 100 µM uridine was selected as the initial concentration to examine the effects of elevated extracellular uridine on islet function. Mouse islets were cultured overnight for 16h in the presence or absence of 100 μM uridine, and uridine was maintained throughout the static GSIS assay. Uridine had no effect on insulin secretion under low-glucose (3 mM glucose) but increased insulin secretion approximately twofold at stimulatory glucose (16.7 mM glucose) (**Fig. 1a**). This insulinotropic effect is highly specific to uridine. Neither uracil, the nucleobase of uridine, nor pseudouridine, a naturally occurring uridine isomer generated by conversion of the C-N glycosidic bond to a C-C bond while retaining the same molecular formula and mass [35], enhanced GSIS (**Fig. 1a,b**). Likewise, uridine monophosphate (UMP), a commonly used dietary supplement for increasing circulating uridine levels [4], failed to potentiate GSIS (**Fig. 1c**). Together, these findings demonstrate that the potentiation of insulin secretion is a unique property of uridine rather than a general feature of structurally related pyrimidine metabolites.

**Figure 1.**
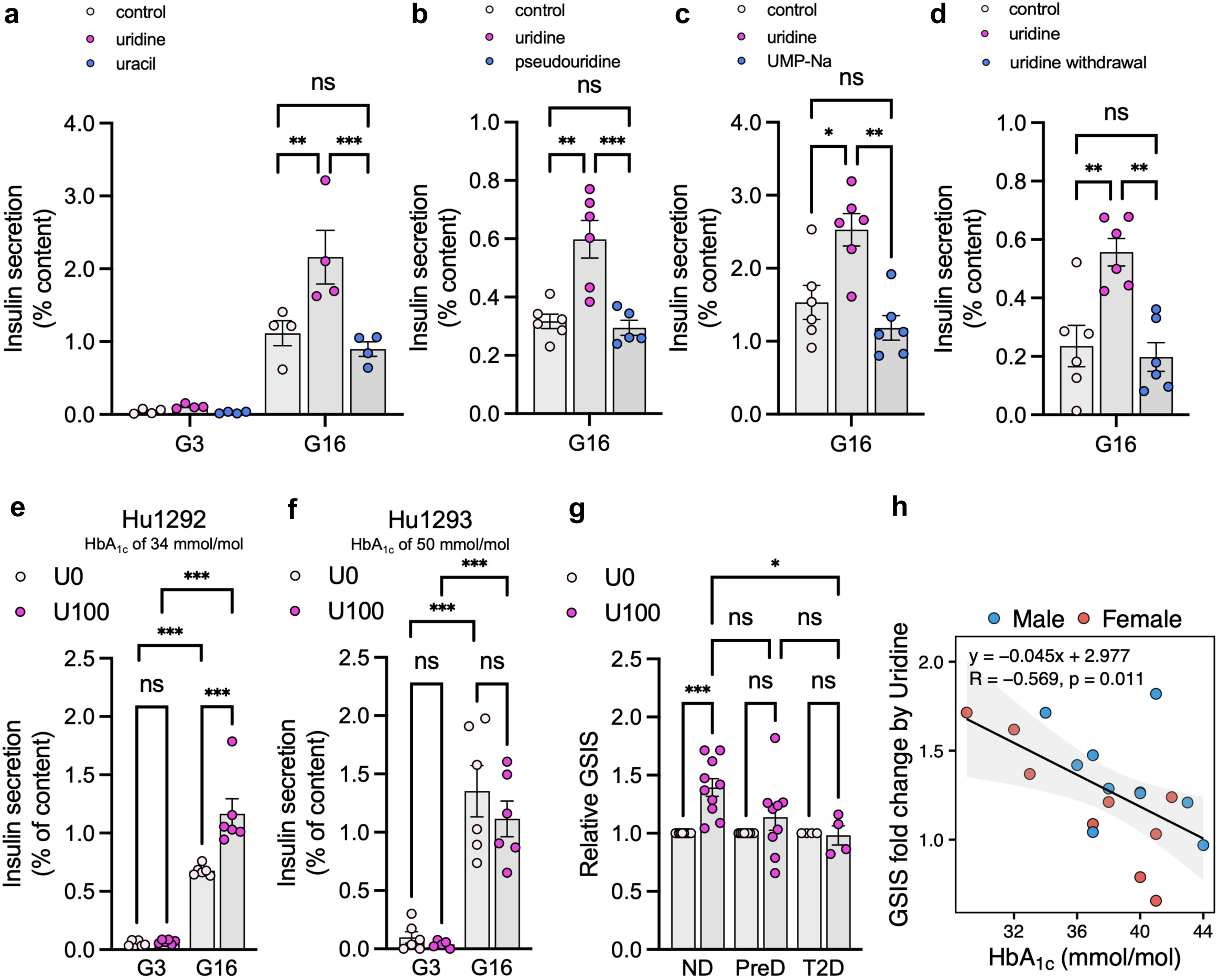
Uridine potentiates glucose-stimulated insulin secretion in mouse and human islets. (**a-d**) Static glucose-stimulated insulin secretion (GSIS) from mouse islets cultured overnight (16 h) with the indicated treatments. Uridine, uracil, pseudouridine, and UMP-Na (100 µM each) were added immediately after islet isolation and maintained throughout the overnight culture and GSIS assay. Control islets received no treatment. Insulin secretion was measured at 3 mM (G3) and 16.7 mM (G16) glucose. **(d)** Reversibility of uridine-induced GSIS potentiation. Mouse islets were cultured with 100 μM uridine for 16 h and either analyzed immediately by GSIS or cultured for an additional 24 h in standard RPMI 1640 medium before GSIS assessment. Data are expressed as mean ± SEM of insulin secretion normalised to the insulin content of 10 islets. Islet insulin content corresponding to panels A–D is shown in ESM Fig. 1a–d. *n* = 4 mouse biological replicates in (**a**), and n = 6 mouse biological replicates in (**b-d**). **(e,f)** Representative static GSIS from human islets isolated from a non-diabetic donor (Hu1292, HbA_1c_ of 34 mmol/mol) and a donor with T2D (Hu1293, HbA_1c_ of 50 mmol/mol). *n* = 6 technical replicates. (**g**) Relative GSIS in untreated (U0) and uridine-treated (U100) human islets. For each donor, glucose-stimulated insulin secretion from untreated islets (U0) was normalised to 1, and insulin secretion from uridine-treated islets (U100) is represented relative to its paired untreated control. Each dot represents one donor (*n* = 10 non-diabetic [ND], *n* = 9 prediabetic [PreD], and *n* = 4 T2D). **(h)** Correlation between donor HbA1c and uridine-induced GSIS potentiation. GSIS fold change by uridine was the relative GSIS in uridine-treated islets. Donor characteristics are provided in ESM Table 1. Data are presented as mean ± SEM. Statistical significance in (**a-g**) was determined by two-way ANOVA followed by Šídák’s multiple-comparison test. Correlation analysis in (**h**) was performed by linear regression. \**p* < 0.05, \*\**p* < 0.01, \*\*\**p* < 0.001; ns, not significant.

To determine whether the enhanced GSIS reflected irreversible islet injury following uridine exposure, a third experimental group was included in which islets were cultured with uridine for 24 h and subsequently allowed to recover in standard culture medium for an additional 24 h before GSIS analysis. Insulin secretion in the withdrawal group was indistinguishable from that of untreated controls (**Fig. 1d**), indicating that uridine-induced potentiation is fully reversible and does not cause persistent impairment of islet function under these experimental conditions. Furthermore, islet insulin content was unchanged across all static GSIS experiments (**ESM Fig. 1a-d**), indicating that the enhanced insulin secretion cannot be explained solely by altered insulin biosynthesis or storage.

Uridine-mediated potentiation of GSIS was also observed in human islets from donors without diabetes but was progressively attenuated in islets from donors with prediabetes or T2D (**Fig. 1e–g**). Across individual donors, the uridine-induced fold increase in GSIS correlated inversely with donor HbA_1c_ (**Fig. 1h; ESM Table 1**), suggesting that worsening glycemic control is associated with reduced islet responsiveness to uridine. In contrast, uridine-mediated GSIS potentiation was not correlated with donor BMI (**ESM Fig. 1e**), indicating that obesity alone was not associated with diminished uridine responsiveness in this donor cohort.

### Uridine-induced potentiation of GSIS is through UPP1-dependent intracellular metabolism

To determine whether uridine acts intracellularly, mouse islets were treated with the equilibrative nucleoside transporter inhibitor dilazep dihydrochloride (DDC). DDC inhibited uridine-induced potentiation of GSIS in a dose-dependent manner without affecting islet insulin content (**Fig**. **2a,b**), indicating that cellular uptake of uridine is required for its insulinotropic effect.

UPP1 is the rate-limiting enzyme responsible for uridine catabolism [14]. To investigate whether UPP1 is necessary for uridine-potentiated GSIS, we examined islets from whole-body *Upp1* knockout (KO) mice, a model previously established for studies of uridine homeostasis [15]. Successful deletion of *Upp1* was confirmed by PCR as described [16] (**ESM Fig. 2a**). Basal and glucose-stimulated insulin secretion under standard culture conditions were comparable between WT and *Upp1* KO islets (**Fig. 2c**). However, unlike WT islets, *Upp1*-deficient islets failed to exhibit uridine-potentiated GSIS (**Fig. 2c**), despite having normal insulin content (**Fig. 2d**).

To determine whether this requirement is through β-cell UPP1, we engineered an inducible β-cell-specific *Upp1* knockout mouse using a Tet-on strategy consisting of an *Ins1* promoter-driven tetracycline transactivator (MIPrtTA) as reported [36] and tetracycline-responsive Cre recombinase (TRE-Cre) (**Fig. 2e**). Tissue-specific recombination was confirmed by PCR using primers specific for the knockout allele; following doxycycline induction, recombination was detected exclusively in islets and not in liver (**ESM Fig. 2b-d**). Consistent with successful gene deletion, UPP1 protein expression was reduced in β-cell-specific *Upp1* knockout (*Upp1* KO^β^) islets as determined by immunoblotting and immunofluorescence (**Fig. 2f,g**). Similar to whole-body *Upp1* knockout islets, uridine-induced potentiation of GSIS was completely abolished in *Upp1* KO^β^ islets, whereas islet insulin contents remained unchanged (**Fig. 1h-i**). Importantly, under standard culture conditions without uridine supplementation, WT and *Upp1* KO^β^ islets exhibited comparable GSIS, indicating that loss of UPP1 does not intrinsically impair β-cell secretory function but specifically eliminates the response to elevated extracellular uridine.

### Uridine potentiation involves transient activation of MEK/ERK signaling pathway

Uridine catabolism by UPP1 has been shown to activate the mitogen-activated protein kinase kinase (MEK)/extracellular signal-regulated kinase (ERK) signaling pathway in the human pancreatic ductal adenocarcinoma cell line ASPC1 under glucose-deprived conditions [34]. Meanwhile, MEK/ERK signaling is known to be activated by glucose and other agents that induce and potentiate insulin secretion [37]. We therefore investigated whether uridine and UPP1 regulate MEK/ERK signaling in pancreatic islets during glucose stimulation. Following equilibration in KRBH containing 3 mM glucose, WT islets were stimulated with 16.7 mM glucose, as in a standard GSIS assay. Glucose stimulation elicited a rapid but transient increase in ERK phosphorylation, with p-ERK peaking at 5 min (**Fig. 3a**, lane 2 versus 1). Uridine treatment markedly enhanced ERK phosphorylation at this 5-min peak (**Fig. 3a**, lane 6 versus 2), while the subsequent decline in p-ERK was unaffected (**Fig. 3a**, lanes 7–8 versus 3–4), indicating that uridine amplified the magnitude, but not the duration, of ERK activation.

**Figure 2.**
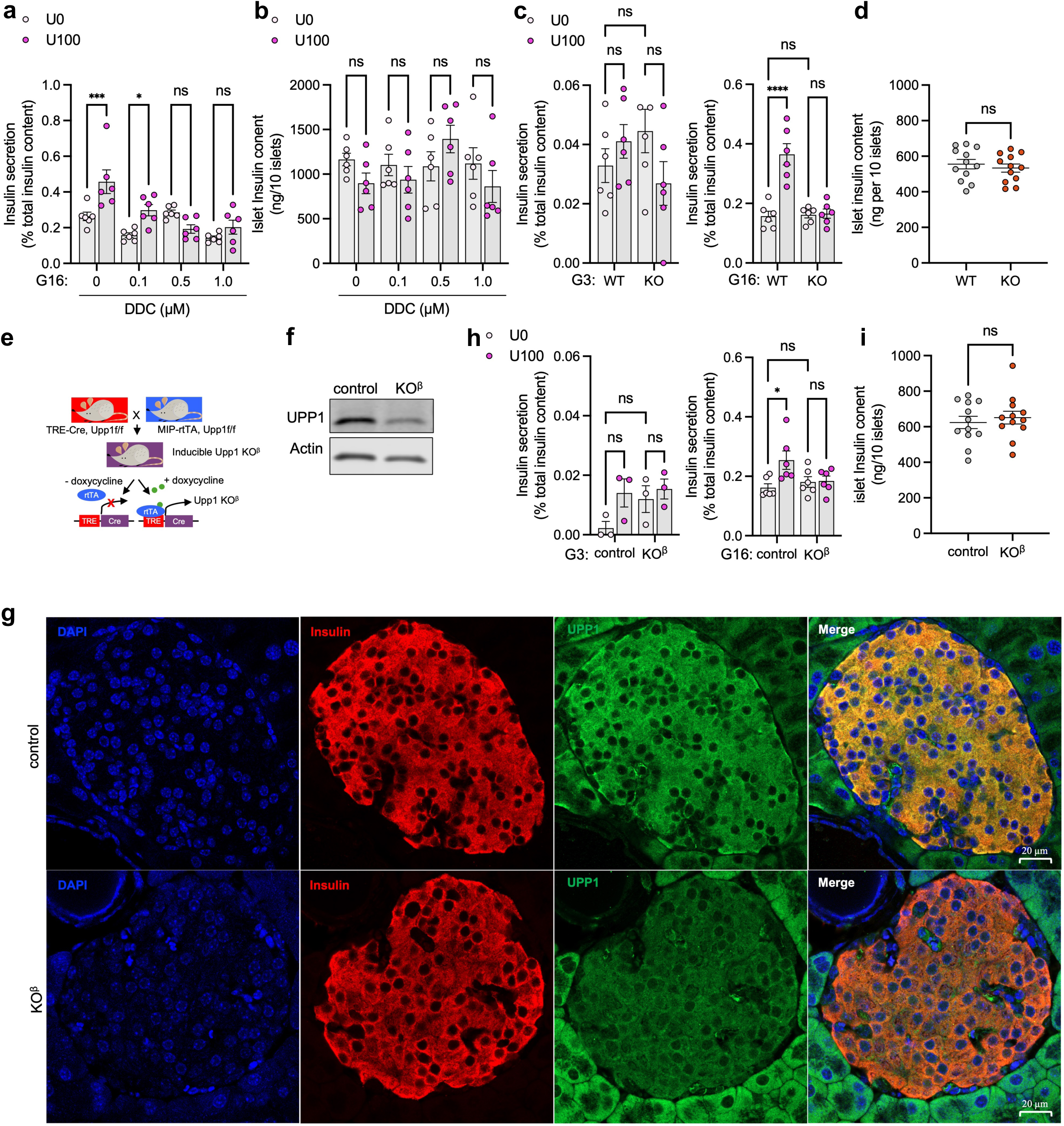
Uridine-mediated potentiation of GSIS requires UPP1. (**a,b**) Static GSIS assays in WT mouse islets treated with dilazep dihydrochloride (DCC). At each indicated DCC concentration, islets were divided into two groups and cultured overnight (16 h) with either 100 μM uridine (U100) or no uridine (U0). Treatments were initiated immediately after islet isolation and maintained throughout the overnight culture and GSIS assay. Insulin secretion was measured at 16.7 mM (G16) glucose, expressed as mean ± SE normalised to the insulin content of 10 islets. *n* = 6 mouse biological replicates. (**c,d**) Static GSIS assays in WT and whole-body *Upp1* KO islets. Islets isolated from each mouse were divided into two groups and cultured overnight with either 100 μM uridine (U100) or no uridine (U0). Insulin secretion was measured at 3 mM (G3) and 16.7 mM (G16) glucose. Insulin secretion is expressed as mean ± SE normalised to the insulin content of 10 islets. *n* = 6 mouse biological replicates per group. (**e**) Breeding strategy and doxycycline induction protocol for inducible β-cell-specific *Upp1* knockout (*Upp1* KO^β^) mice.**(f)** Representative immunoblot of islet UPP1 protein expression in 10-week-old male control (*Upp1*^f/f^, MIPrtTA) and *Upp1* KO^β^ (*Upp1*^f/f^, MIPrtTA, TRE-Cre) mice following 3 weeks of Dox600 chow feeding. (**g**) Representative immunofluorescence images of pancreatic sections from male control and *Upp1* KO^β^ mice co-stained for insulin (Thermo 14-9769-82) and UPP1 (Proteintech, 14186-1-AP). Images were acquired using a Zeiss LSM900 confocal microscope with a 63x objective. Scale bar, 20 μm. Mice were analyzed after 10 weeks of Dox600 chow feeding. **(h,i)** Static GSIS assays in control and *Upp1* KO^β^ islets. Mice were fed Dox600 chow for 7 weeks before islet isolation. Islets from each mouse were divided into two groups and cultured overnight with either 100 μM uridine (U100) or no uridine (U0). Insulin secretion is expressed as mean ± SE and normalised to the insulin content of 10 islets. *n* = 3 mouse biological replicates per group for basal insulin secretion (G3) and *n* = 6 mouse biological replicates per group for glucose-stimulated insulin secretion (G16). Data are presented as mean ± SEM. Statistical significance in (**a-d**) and (**h,i**) was determined by two-way ANOVA followed by Šídák’s multiple-comparison test. \**p* < 0.05, \*\**p* < 0.01, \*\*\**p* < 0.001; ns, not significant.

**Figure 3.**
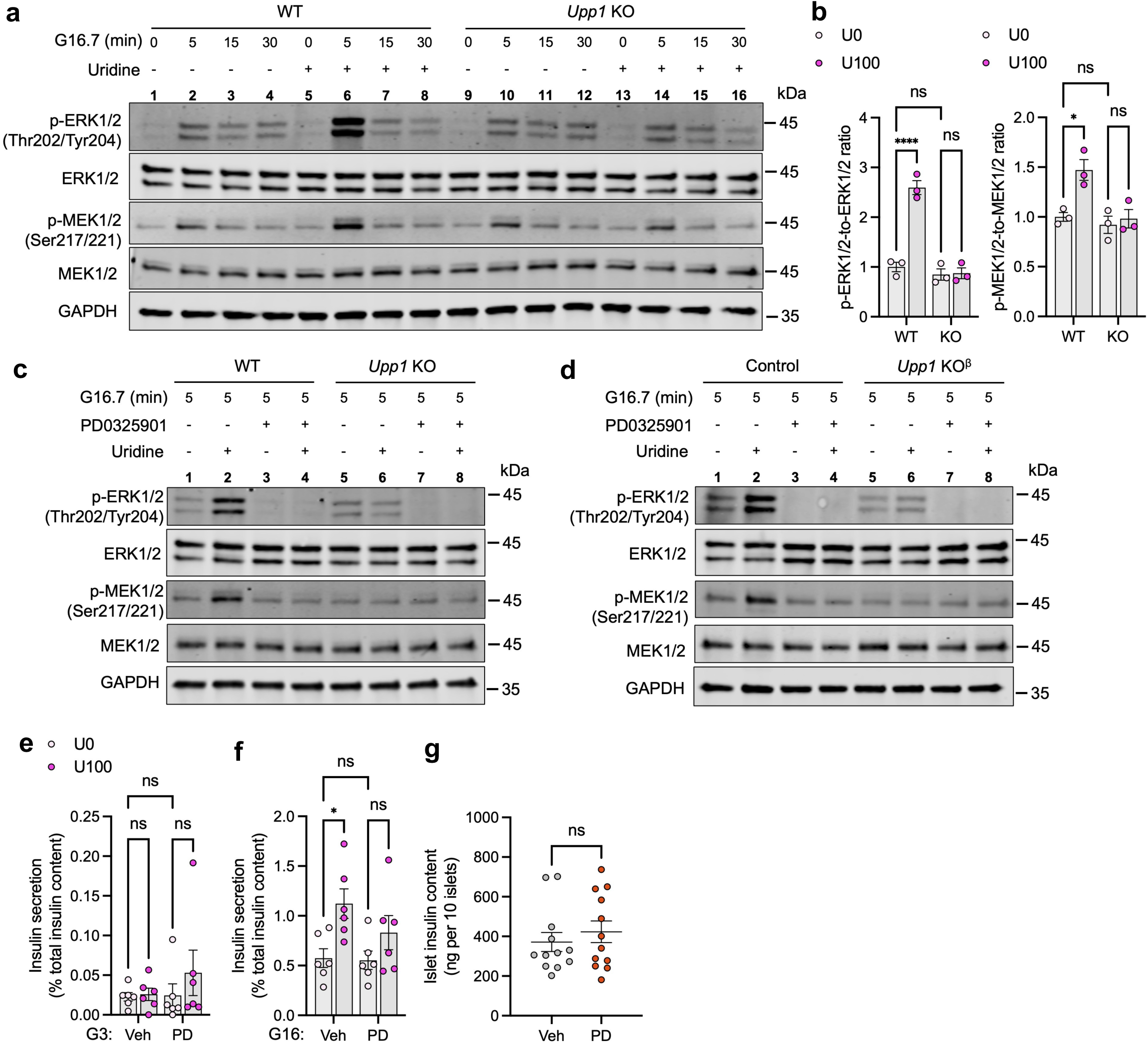
Uridine potentiation of glucose-stimulated insulin secretion requires transient activation of the MAPK pathway. (**a**) WT and *Upp1* KO islets from each mouse were divided into two groups: untreated (U0) and uridine-treated (U100). Uridine treatment was initiated during overnight culture immediately after islet isolation and maintained throughout the GSIS assay. Following a 2-h acclimation in KRBH buffer containing 3 mM glucose, islets were stimulated with 16.7 mM glucose for 0, 5, 15, or 30 min. Islets were then lysed and analyzed by SDS-PAGE and immunoblotting for phosphorylated ERK1/2 (p-ERK1/2) and total ERK1/2, as well as phosphorylated MEK1/2 (p-MEK1/2) and total MEK1/2. (**b**) Quantification of the p-ERK1/2-to-total ERK1/2 ratio at 5 min after glucose stimulation from the experiments shown in (**a**). The average p-ERK1/2-to-total ERK1/2 ratio in untreated WT islets within each independent experiment was normalised to 1. *n* = 3 independent experiments, with each experiment using islets pooled from 4 mice per genotype. (**c**) The uridine-induced enhancement of MAPK signaling at 5 min after glucose stimulation was abolished in *Upp1* KO islets. Treatment with the MEK1/2 inhibitor PD0325901 (PD) eliminated ERK phosphorylation in response to glucose stimulation, regardless of uridine treatment. Two independent experiments were performed, with each experiment using islets pooled from four mice per genotype. (**d**) The uridine-induced enhancement of MAPK signaling at 5 min after glucose stimulation was also abolished in *Upp1* KO^β^ islets. PD0325901 inhibited ERK phosphorylation irrespective of uridine treatment. Control (*Upp1* f/f, MIPrtTA) and *Upp1* KO^β^ (*Upp1* f/f, MIPrtTA, TRE-Cre) mice were fed Dox600 chow for 7 weeks to induce β-cell-specific deletion of *Upp1* before islet isolation. Two independent experiments were performed, with each experiment using islets pooled from four mice per genotype. (**e-g**) PD0325901 abolished uridine potentiation of GSIS (**f**) without affecting baseline insulin secretion (**e**) or islet insulin content (**g**). Insulin secretion is expressed as mean ± SEM and normalised to the insulin content of 10 islets. *n* = 6 mouse biological replicates. Data are presented as mean ± SE. Statistical significance in (**b**), (**e**) and (**f**) was determined by two-way ANOVA followed by Šídák’s multiple-comparison test, whereas (**g**) was analyzed using a two-tailed unpaired Student’s t-test. \**p* < 0.05, \*\*\**p* < 0.001; ns, not significant.

*Upp1* KO islets also exhibited a transient increase in ERK phosphorylation 5 min after glucose stimulation (**Fig. 3a**, lane 10 versus 9). However, unlike WT islets, uridine failed to further enhance ERK phosphorylation in *Upp1* KO islets (**Fig. 3a**, lane 14 versus 10). Quantification of the p-ERK/ERK ratio showed that uridine increased ERK phosphorylation by approximately 2.6-fold in WT islets at the 5-min time point, whereas no significant effect of uridine was observed in *Upp1* KO islets (**Fig. 3b**).

Phosphorylation of MEK closely paralleled that of ERK in both WT and *Upp1* KO islets with respect to the magnitude, timing, and transient nature of the response (**Fig. 3a,b**). Together, these findings demonstrate that uridine potentiates glucose-induced activation of the MEK/ERK signaling pathway in a UPP1-dependent manner.

The 0-min time point in **Fig**. **3a** corresponds to the end of the 2-h acclimation period in KRBH containing 3 mM glucose. The absence of increased ERK phosphorylation in uridine-treated islets at this time point indicates that uridine alone does not activate MEK/ERK signaling under low-glucose conditions (**Fig. 3a**, lanes 5 and 13 versus 1 and 9). Likewise, WT and *Upp1* KO islets exhibited comparable levels of MEK/ERK phosphorylation following overnight culture in RPMI 1640 medium containing 8 mM glucose (**ESM Fig. 3**), demonstrating that UPP1 deficiency does not alter basal MAPK signaling under standard culture conditions. Furthermore, overnight uridine treatment did not affect MEK or ERK phosphorylation in either WT or *UPP1* KO islets (**ESM Fig. 3**), demonstrating that uridine does not modulate basal MAPK signaling irrespective of UPP1 expression. Consistent with these findings, islets isolated from *Upp1* KO^β^ mice exhibited MEK and ERK phosphorylation levels comparable to those of control islets, regardless of uridine supplementation (**ESM Fig. 4**). Together, these results indicate that neither uridine nor UPP1 deficiency affects basal MEK/ERK signaling and that UPP1-dependent activation of the MAPK pathway occurs specifically in response to glucose stimulation.

**Figure 4.**
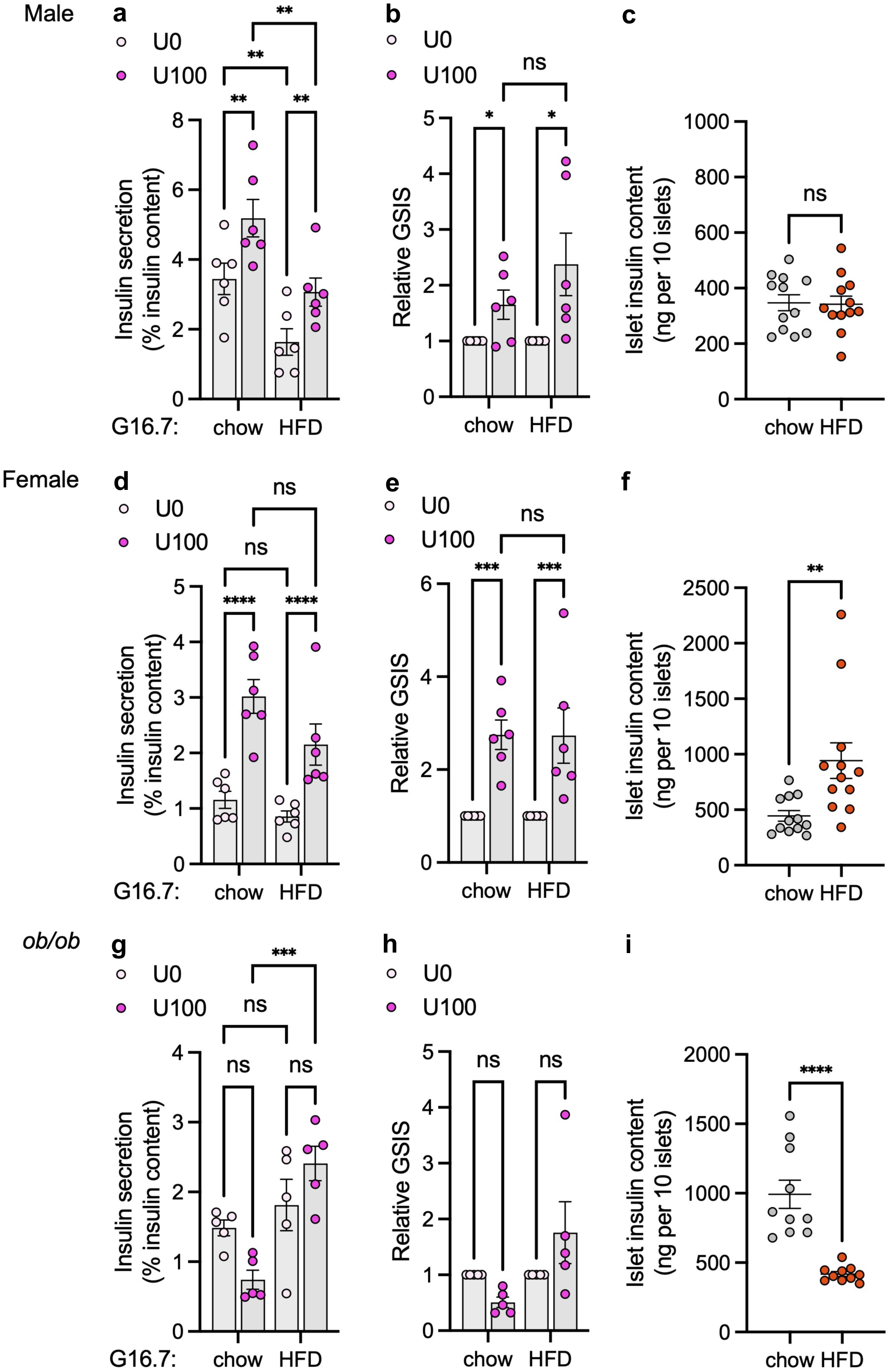
Uridine potentiation of GSIS is preserved in HFD-fed mice but abolished in *ob/ob* mice. WT mice were switched to a HFD at 10 weeks of age and maintained on HFD for 20 weeks before islet isolation. Age-matched chow-fed mice were used as controls, and islets from chow– and HFD-fed mice were isolated and assayed on the same day. For each mouse, isolated islets were divided into two groups: untreated (U0) and uridine-treated (U100). Uridine treatment was initiated during overnight culture immediately after islet isolation and maintained throughout the GSIS assay. (**a, d, g**) Insulin secretion was measured at 16.7 mM (G16.7) glucose. Glucose-stimulated insulin secretion is expressed as mean ± SEM and normalised to the insulin content of 10 islets. *n* = 6 biological replicates per group for WT chow– and HFD-fed mice and *n* = 5 mouse biological replicates per group for chow– and HFD-fed *ob/ob* mice. (**b, e, h**) Relative GSIS in untreated (U0) and uridine-treated (U100) islets. For each mouse, glucose-stimulated insulin secretion from untreated islets (U0) was normalised to 1, and insulin secretion from uridine-treated islets (U100) is represented relative to its paired untreated control. (**c, f, i**) Islet insulin contents used to normalize GSIS. Data are presented as mean ± SEM. Statistical significance in **a, b**, **d**, **e**, **g**, and **h** was determined by two-way ANOVA followed by Šídák’s multiple-comparison test. **c**, **f**, and **i** were analyzed using two-tailed unpaired Student’s t-test. \**p* < 0.05, \*\**p* < 0.01, \*\*\**p* < 0.001; ns, not significant.

To examine whether activation of MEK/ERK pathway is required for uridine potentiation of GSIS, islets were treated with the selective MEK1/2 inhibitor PD0325901, as described previously [24], throughout the overnight culture period and the subsequent glucose stimulation in KRBH buffer. PD0325901 completely abolished glucose-stimulated ERK phosphorylation in WT and *Upp1* KO islets (**Fig. 3c**, lanes 1 and 3 for WT, lanes 5 and 7 for *Upp1* KO). Likewise, PD0325901 eliminated ERK phosphorylation in islets treated with uridine (**Fig. 3c**, lanes 2 and 4 for WT, lanes 6 and 8 for *Upp1* KO), indicating that both glucose– and uridine-induced ERK phosphorylation are abolished by the MEK1/2 inhibitor. Notably, PD0325901 also abolished the uridine-induced increase in MEK phosphorylation in WT islets, while having little effect on glucose-induced MEK phosphorylation (**Fig. 3c**, lane 4 versus 2, and 3 versus 1), suggesting that unlike glucose-induced MEK activation, uridine-induced MEK phosphorylation is dependent on ERK activity, consistent with a positive feedback mechanism within the MEK/ERK signaling cascade. Similar results were observed in islets from *Upp1* KO^β^ mice (**Fig. 3d**).

We next examined the impact of PD0325901 on GSIS from WT islets. As expected, PD0325901 completely abolished the uridine-induced enhancement of GSIS without altering baseline insulin secretion (**Fig. 3e,f**). Importantly, PD0325901 did not alter islet insulin contents (**Fig. 3g**), indicating that inhibition of ERK signaling during the course of the experiment did not impair β-cell insulin biosynthesis or storage. Thus, the loss of uridine potentiation following PD0325901 treatment is unlikely secondary to β-cell exhaustion. To exclude the possibility that prolonged exposure to PD0325901 contributed to this effect, the inhibitor was applied only during the final 2 h of overnight culture in RPMI 1640 medium before the islets were transferred to KRBH. Under these conditions, uridine-potentiated GSIS was still completely abolished (**ESM Fig. 5**), indicating that acute inhibition of ERK signaling is sufficient to block the insulinotropic effect of uridine. Notably, inhibition of ERK phosphorylation had no effect on GSIS in the absence of uridine (**Fig. 3e, ESM Fig. 5a**), consistent with a previous report demonstrating that glucose-induced MAPK activation is dispensable for GSIS in the INS-1 β-cell line [38]. Together, these findings indicate that MEK/ERK signaling is specifically required for uridine-mediated potentiation of insulin secretion but is not essential for basal glucose-stimulated insulin secretion.

**Figure 5.**
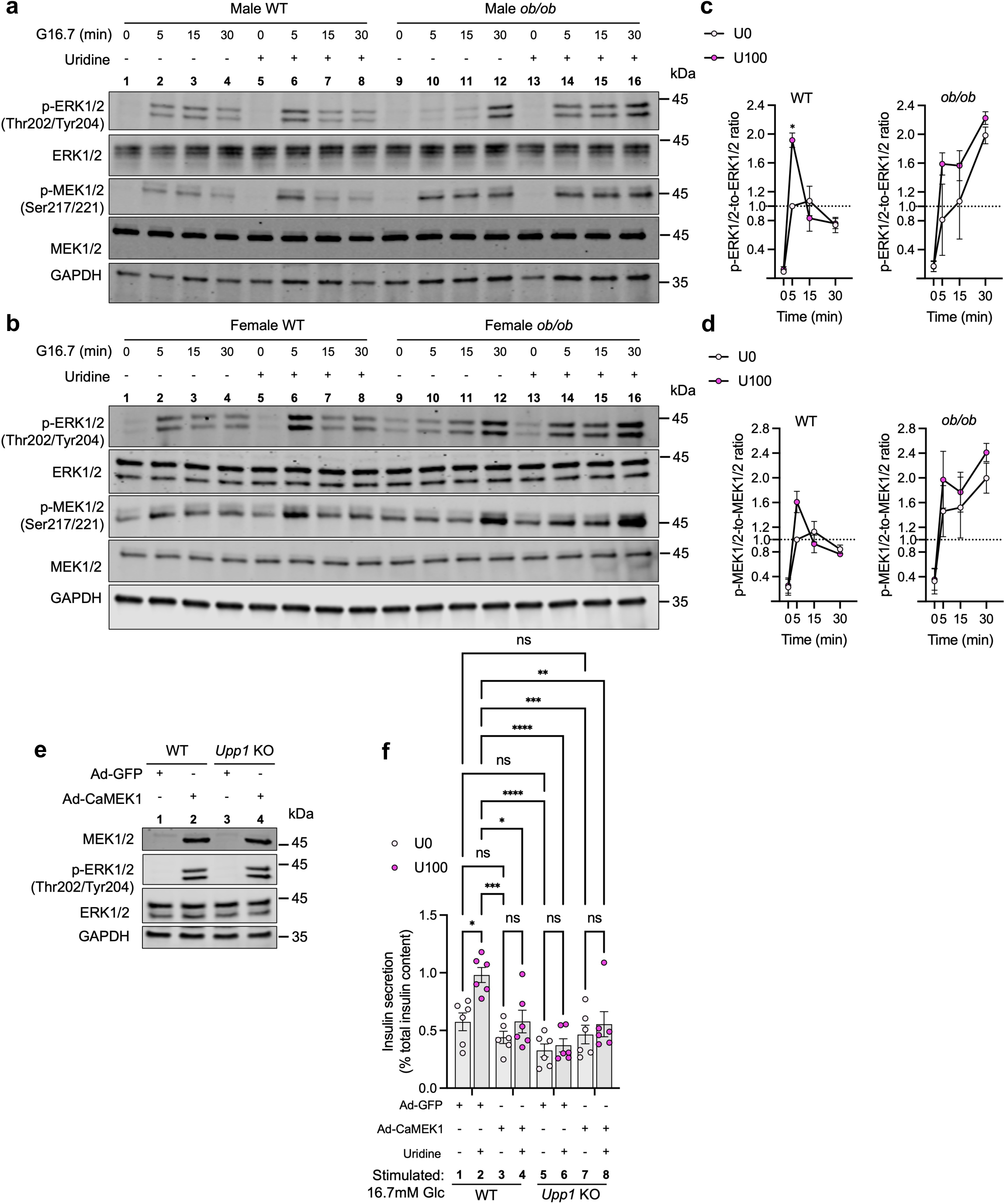
Aberrant MAPK signaling dynamics in islets from *ob/ob* mice and the requirement for transient ERK activation in uridine. (**a,b**) Islets from each mouse were divided into two groups: untreated (U0) and treated (U100). Uridine treatment began with overnight incubation after islet isolation and continued throughout the entire GSIS assay. Mouse islets were incubated in KRBH containing 3 mM glucose for 2 h, then stimulated with 16.7 mM glucose for 0, 5, 15, or 30 min. Islets were lysed for SDA-PAGE and immunoblotting for phosphorylated ERK1/2 (p-ERK) and total ERK1/2, as well as phosphorylated MEK1/2 (p-MEK1/2) and total MEK1/2. (**c,d**) Quantification of the p-ERK1/2-to-total ERK1/2 ratio (**c**) and the p-MEK1/2-total MEK1/2 ratio (**d**) from the experiments shown in (**a,b**) plus another biological replicate. The p-ERK1/2-to-total ERK1/2 ratio of untreated WT islets at 5 min after glucose stimulation was normalised to 1 within each independent experiment. *n* = 3 experimental replicates, with each replicate comprising islets pooled from 2 *ob/ob* mice and, separately, eight WT mice. (**e**) WT and *Upp1* KO islets were transduced with adenovirus expressing GFP or constitutively active MEK1 (CaMEK1). Forty-eight hours after transduction in RPMI 1640 media containing 8 mM glucose, islets were analyzed by immunoblotting for CaMEK1 expression and ERK1/2 phosphorylation. The immunoblot shown was selected to optimally visualize CaMEK1 overexpressed in lanes 2 and 4 and the corresponding increase in p-ERK1/2. Because of the high abundance of CaMEK1 protein, endogenous MEK1/2 and p-ERK1/2 bands in lanes 1 and 3 are less apparent than those shown in ESM Fig. 3a. (**f**) Adenovirus-transduced WT and *Upp1* KO islets were subjected to GSIS assays. Insulin secretion is expressed as mean ± SEM and normalised to the insulin content of 10 islets. *n* = 6 mouse biological replicates per group. Data are presented as mean ± SE. Statistical significance in **c**, **d** and **f** was determined by two-way ANOVA followed by Šídák’s multiple-comparison test. \**p* < 0.05, \*\**p* < 0.01, \*\*\**p* < 0.001; ns, not significant.

### Uridine potentiation of GSIS is abolished in *ob/ob* mice but preserved in HFD-fed mice

Because obesity is associated with disrupted uridine homeostasis, we next examined whether uridine potentiation of GSIS is altered in islets from HFD-fed and leptin-deficient *ob/ob* mice. In male mice, islets from HFD-fed animals exhibited lower GSIS than those from chow-fed controls, regardless of uridine treatment (**Fig. 4a**). To directly compare the potentiation effect of uridine independent of basal GSIS, the relative GSIS was calculated for each islet preparation as previously described for human islets. Uridine enhanced GSIS by 1.65-fold in chow-fed mice and by 2.37-fold in HFD-fed mice (**Fig. 4b**), indicating that HFD feeding did not impair the uridine potentiation. Islet insulin content was comparable between the two groups (**Fig. 4c**).

In female mice, HFD feeding did not significantly alter GSIS compared with chow-fed controls, either in the presence or absence of uridine (**Fig. 4d**). Uridine increased GSIS by 2.75-fold in chow-fed mice and 2.73-fold in HFD-fed mice (**Fig. 4e**), demonstrating that uridine potentiation was fully preserved in female HFD-fed mice. Although islet insulin content was significantly higher in HFD-fed females than in chow-fed controls (**Fig. 4f**), this difference did not affect the magnitude of uridine potentiation under each feeding condition. Together, these findings indicate that diet-induced obesity does not diminish the ability of uridine to potentiate GSIS in isolated islets.

In contrast, islets from *ob/ob* mice failed to respond to uridine. GSIS was comparable between uridine-treated and untreated islets from *ob/ob* mice under both chow– and HFD-fed conditions (**Fig. 4g,h**), indicating a complete loss of uridine potentiation. Islet insulin content was significantly lower in HFD-fed *ob/ob* mice than in chow-fed *ob/ob* mice (**Fig. 4i**). Collectively, these results demonstrate that uridine potentiation of GSIS is selectively abolished in leptin-deficient obesity but remains intact in diet-induced obesity.

### Mitogen-activated protein kinase (MAPK) signaling dynamics and UPP1 expression are altered in islets from *ob/ob* mice

Unlike WT islets, islets from *ob/ob* mice failed to exhibit the characteristic transient peak of ERK phosphorylation 5 min after glucose stimulation (**Fig. 5a,b**, lane 10 versus 2). Instead, ERK phosphorylation increased progressively throughout the first 30 min of glucose stimulation without returning toward baseline (**Fig. 5a,b**, lanes 10–12 for *ob/ob* versus 2–4 for WT). Uridine treatment further enhanced ERK phosphorylation at the 5-min time point in *ob/ob* islets; however, ERK phosphorylation remained persistently elevated through 30 min rather than declining as observed in WT islets (**Fig. 5a,b**, lanes 14–16 for *ob/ob* versus 6–8 for WT islets). Consequently, glucose– and uridine-induced MAPK signaling in *ob/ob* islets displayed a markedly different temporal patterns from that observed in either WT or *Upp1 KO* islets (**Fig. 5c**). Consistent with ERK phenotype, the transient increase in MEK phosphorylation normally induced by glucose stimulation was also disrupted in *ob/ob* islets (**Fig. 5d**).

To determine whether the transient nature of ERK activation is required for uridine potentiation of GSIS, we introduced constitutively active MEK1 (CaMEK1), as previously described [25, 26], into WT and *Upp1* KO islet cells using adenoviral transduction. CaMEK1 increases ERK phosphorylation in both WT and *Upp1* KO islets cultured in RPMI 1640 media containing 8 mM glucose (**Fig. 5e**). However, constitutive MEK activation failed to restore uridine potentiation in *Upp1* KO islets (**Fig. 5f**, lanes 7 and 8). Moreover, persistent activation of the MEK/ERK pathway abolished uridine-potentiated GSIS in WT islets (**Fig. 5f**, lanes 3 and 4). Importantly, constitutive MEK activation did not alter GSIS in the absence of uridine (**Fig. 5f**, lanes 1 and 3), suggesting that the MAPK activation is dispensable for glucose-stimulated insulin secretion, consistent with our finding that pharmacological inhibition of MEK/ERK signaling likewise had no effect on GSIS in standard assay (**Fig. 3e**). Together, these results indicate that uridine potentiation requires a precisely regulated, transient activation of the MEK/ERK pathway.

Both failure to initiate this response, as observed in UPP1 deficient islets, and failure to terminate it, as observed in *ob/ob* islets, abolish the insulinotropic effect of uridine.

We next examined whether altered UPP1 expression was associated with the abnormal MAPK signaling observed in *ob/ob* islets. Two anti-UPP1 antibodies were validated for immunoblot detection of UPP1 in islet lysates (**ESM Fig. 6a-e**) and subsequently used for quantitative analyses. UPP1 protein abundance was approximately twofold higher in islets from *ob/ob* mice than in WT controls (**Fig. 6a**), whereas *Upp1* mRNA expression was markedly reduced (**Fig. 6b**), suggesting post-transcriptional accumulation of UPP1 protein. In contrast, UPP1 protein abundance and mRNA expression were unchanged in islets from HFD-fed mice compared with chow-fed controls (**Fig. 6c,d**).

**Figure 6.**
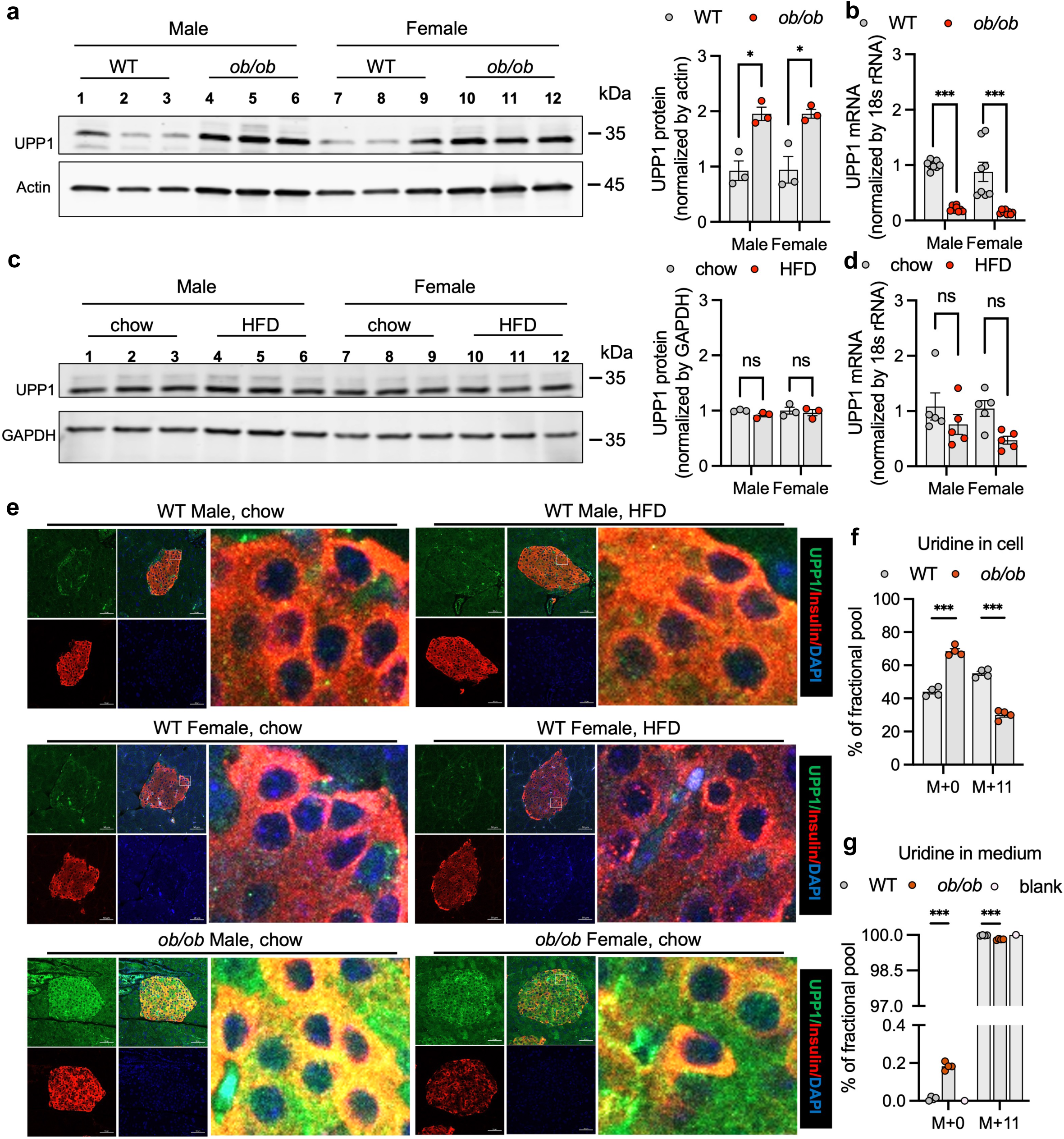
UPP1 expression is increased in islets from *ob/ob* mice but unchanged in HFD-fed mice. (**a,b**) UPP1 protein abundance and mRNA expression in whole-islets isolated from WT and *ob/ob* mice. (**a**) Islet lysate were immunoblotted using an anti-UPP1 antibody (Proteintech). UPP1 protein abundance was normalised by Actin. *n* = 3 mouse biological replicates (**b**) *Upp1* mRNA levels were normalised by 18s rRNA. *n* = 4 mouse biological replicates with 2 technical repeats per mouse. (**c,d**) UPP1 protein abundance and gene expression in whole-islets isolated from mice fed a HFD for 10 weeks and age-matched chow-fed controls. (**c**) Islet lysates were immunoblotted with an anti-UPP1 antibody (LSBio). UPP1 protein abundance was normalised by GAPDH. *n* = 3 mouse biological replicates. (**d**) *Upp1* mRNA levels were normalised by 18s rRNA. *n* = 5 mouse biological replicates. (**e**) Representative immunofluorescence images of pancreatic sections co-stained for insulin and UPP1. Scale bar, 50 μm. (**f,g**) Fractional enrichment of M+0 and M+11 uridine in cell and medium. *n* = 4 experimental replicates for *ob/ob* islets, with each replicate derived from one mouse, and n = 4 experimental replicates for WT islets, with each replicate comprising islets pooled from 4 WT mice. Data are presented as mean ± SEM. Statistical significance in (**a-d)**, and **(f,g)** was determined by two-way ANOVA followed by Šídák’s multiple-comparison test. \**p* < 0.05, \*\*\**p* < 0.001; ns, not significant.

Immunofluorescence staining of pancreatic sections revealed a homogeneous cytosolic distribution of UPP1 protein within β-cells of islets from both male and female mice (**Fig. 6e**, WT male and female, chow), and this pattern was maintained following HFD feeding (**Fig. 6e**, WT male and female, HFD). In contrast, β-cells from *ob/ob* mice exhibited markedly stronger UPP1 staining while retaining its cytosolic localization (**Fig. 6e**, *ob/ob* male and female, chow). These findings demonstrate that leptin deficiency, but not diet-induced obesity, is associated with aberrant accumulation of UPP1 in pancreatic β-cells, providing a potential mechanism for the altered MAPK signaling dynamics and loss of uridine potentiation observed in *ob/ob* islets.

Impaired uridine uptake or excessive uridine catabolism by increased UPP1 could potentially contributed to the loss of uridine responsiveness in *ob/ob* islets. To investigate these possibility, islets were incubated for 1 h in KRBH buffer containing 100 μM [U-^15^N_2_, U-^13^C_9_]uridine and 16.7 mM glucose. Total intracellular uridine abundance was greater in *ob/ob* islets than in WT islets, whereas total medium uridine abundance was comparable between genotypes and remained similar to that in blank medium without islets (**ESM Fig. 6f,g**). Despite the larger intracellular uridine pool, the fractional enrichment of intracellular M+11 uridine was significantly lower in *ob/ob* islets (**Fig. 6f**), indicating reduced contribution of exogenous uridine to the intracellular uridine pool. Conversely, the abundance of unlabeled M+0 uridine in the medium was greater after incubation with *ob/ob* islets (**Fig. 6g**), consistent with increased release of endogenous uridine. Collectively, these findings suggest that *ob/ob* islets exhibit reduced uptake and/or retention of exogenous uridine together with increased release of endogenous uridine. However, because the total intracellular uridine pool was not depleted, limited intracellular uridine availability is unlikely to explain the loss of uridine responsiveness in *ob/ob* islets. These measurements do not directly determine whether increased UPP1 abundance is accompanied by increased uridine catabolic flux.

### Islet UPP1 protein expression positively correlates with glycemic deterioration in human donors

UPP1 protein abundance was quantified in human islets (**Fig. 7a**) and examined in relation to donor HbA_1c_. When male (*n* = 10) and female (*n* = 6) donors–including two male donors with prediabetes–were analyzed together, islet UPP1 protein abundance correlated positively with HbA_1c_ (**Fig. 7b).** This positive relationship remained evident when male and female donors were analyzed separately **(Fig. 7c** and **ESM Table 2**), indicating that islet UPP1 abundance increases with worsening glycemic control in both sexes. In contrast, UPP1 protein abundance was not correlated with donor BMI (**ESM Fig. 7a,b**), suggesting that obesity alone was not associated with increased islet UPP1 abundance in this donor cohort.

**Figure 7.**
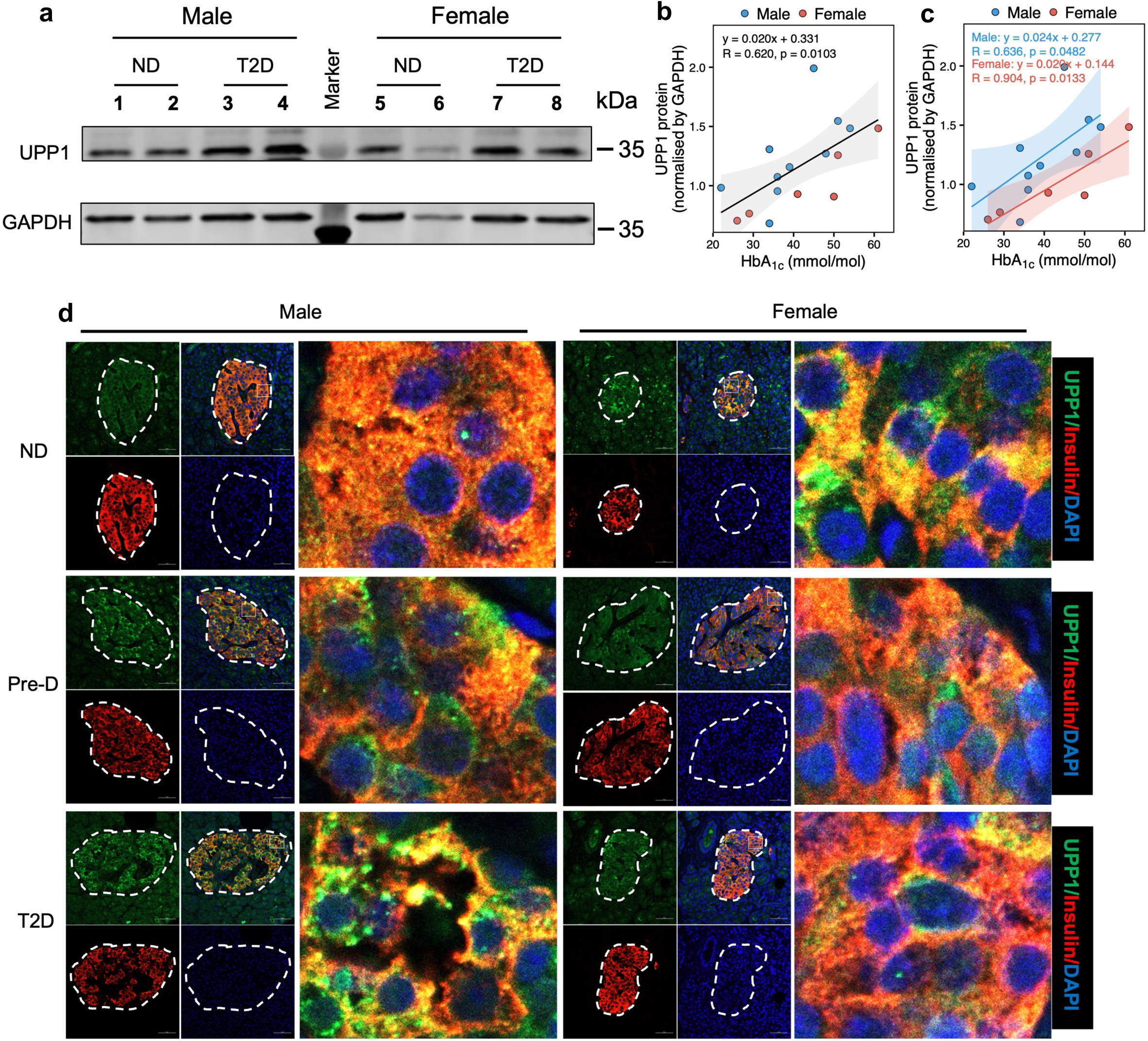
UPP1 expression in pancreatic islets from human donors across the spectrum of glycemic control. (**a**) Representative immunoblot showing UPP1 protein abundance in isolated human islets from donors with varying glycemic status. Islet lysates were immunoblotted with an anti-UPP1 antibody (LSBio) with GAPDH serving as the loading control. (**b,c**) Correlation between islet UPP1 protein abundance and donor HbA_1c_ for all donors combined (**b**) or after stratification by sex (**c**). (**d**) Representative immunofluorescence images of pancreatic sections from male and female donors without diabetes (ND), with prediabetes (Pre-D), or with T2D, co-stained for insulin and UPP1. Scale bar, 50 μm. Data in (**b,c**) were analyzed by linear regression. Donor characteristics are provided in **ESM Table 2** and **3**. Representative immunofluorescence images from each donor are shown in **ESM** Fig. 7c.

We further examined *UPP1* expression during diabetes progression using a human pancreatic islet single-cell RNA-seq dataset (**GSE221156**). β-cell *UPP1* expression was significantly increased in donors with prediabetes relative to donors without diabetes but subsequently decreased in donors with T2D (**ESM Fig. 7c-f**). Moreover, β-cell *UPP1* expression was not correlated with donor HbA_1c_ in this cohort (**ESM Fig. 7g-i**). The discordance between UPP1 protein abundance and transcript expression suggests that *UPP1* may be regulated post-transcriptionally during diabetes progression.

To determine whether the subcellular localization of UPP1 changes during diabetes progression, pancreatic sections from donors without diabetes, with prediabetes, or with T2D (**ESM Table 3** and **ESM Fig. 8**) were immunostained for UPP1. In islets from both male and female donors, UPP1 exhibited either a diffuse or punctate cytosolic distribution, with substantial heterogeneity among individual islets (**Fig. 7d**). No consistent differences in UPP1 subcellular localization were observed between sexes or across stages of diabetes progression, indicating that its heterogeneous intracellular distribution was not clearly associated with sex or glycemic status in this donor cohort.

**Figure 8.**
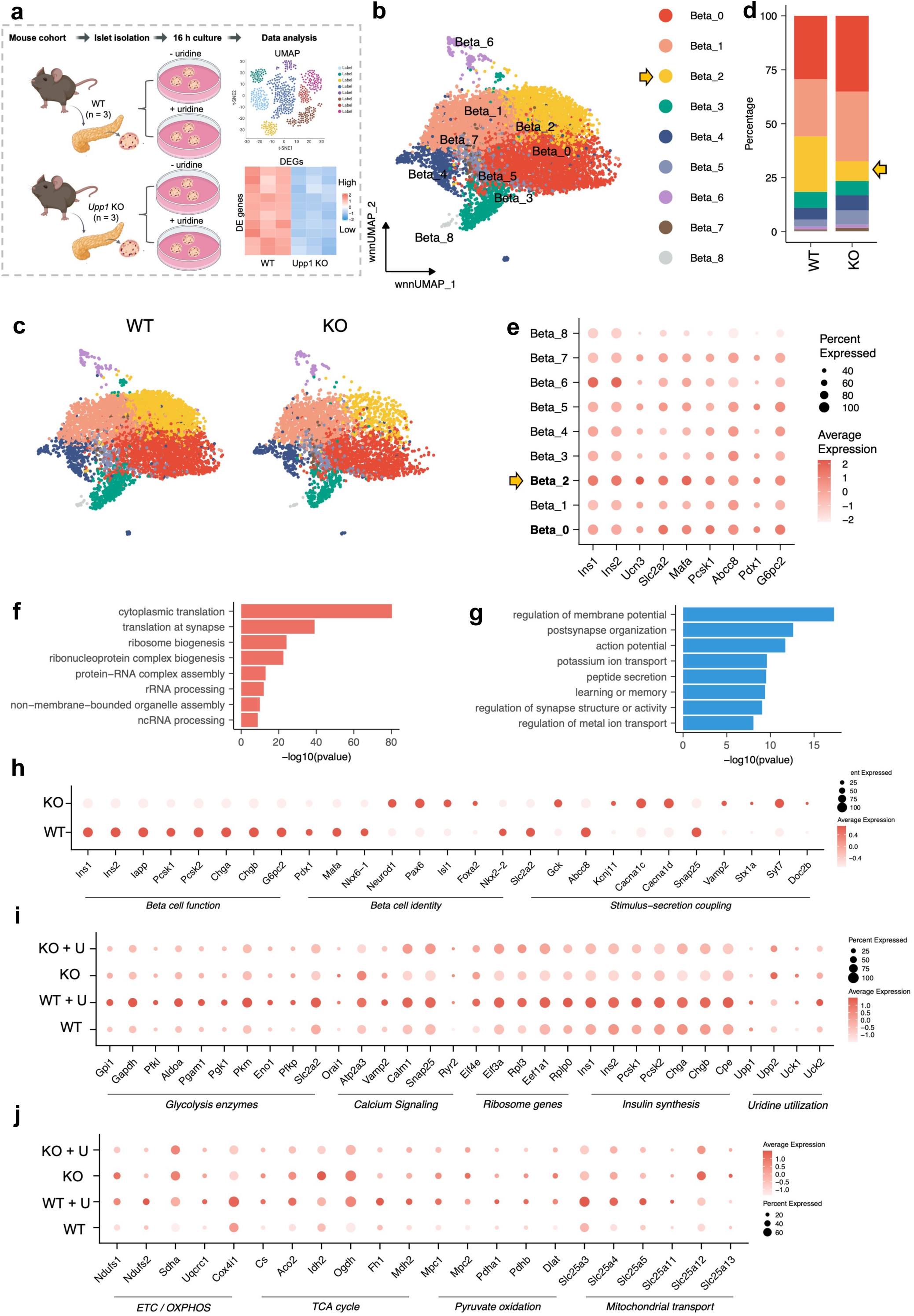
Single-nucleus multiomic analysis of islet β-cells of *Upp1* KO mice. (**a**) Islets were isolated from WT and *Upp1* KO mice, and islets from each mouse were divided equally for overnight culture with or without uridine supplementation. For each genotype and treatment condition, equal numbers of islets from three mice were pooled for subsequent single-nucleus multiomic analysis. Thus, each pooled sample contained islets from three biological donors, with the untreated and uridine-treated samples derived from paired aliquots of the same mice. (**b**) UMAP visualization of β-cell subclusters identified by unsupervised clustering within WT and *Upp1* KO islets. (**c**) Split view of each β-cell subcluster in WT and *Upp1* KO islets on UMAP of **b**. (**d**) Quantification of β-cell subcluster abundance showing a significant reduction in the Beta_2 subcluster in *Upp1* KO islets. (**e**) Dot plot showing the expression of key β-cell identity and functional genes in β-cell subclusters. (**f,g**) Gene Ontology enrichment analysis of top upregulated and downregulated pathways that are enriched in the Beta_2 subcluster. (**h**) Dot plot showing pseudo-bulk differential gene expression of β-cells comparing WT and *Upp1* KO islets. Shown are selected genes involved in β-cell function, identity, and stimulated secretion coupling. (**I,j**) Dot plots showing the transcriptional response to overnight uridine treatment in WT and *Upp1* KO β-cells by pseudo-bulk analysis. Uridine induced coordinated upregulation of genes involved in glycolysis, calcium signaling, ribosome biogenesis, insulin biosynthesis, oxidative phosphorylation, TCA cycle, pyruvate oxidation, and mitochondrial metabolite transport in WT islets, whereas these responses were markedly attenuated or absent in *Upp1* KO islets. *Upp1* expression was increased following uridine treatment in WT β-cells. Data are derived from integrated single-nucleus multiomic (snRNA-seq/snATAC-seq) analyses of isolated mouse islets. Statistical methods and sample sizes are provided in the Methods and **ESM Table 4**. Additional details regarding the experimental design and its limitations are provided in **ESM** Fig. 9a.

### UPP1 is required for maintenance of the β-cell transcriptional program and uridine-induced transcriptional remodeling

Mouse islets from WT and *Upp1* KO mice were subjected to integrated single-nucleus multiomic (snRNA-seq/snATAC-seq) analyses (**Fig. 8a, ESM Fig. 9a, ESM Table 4**). Islet cell composition was comparable between WT and *Upp1* KO mice (**ESM Fig. 9b**). However, analysis of β-cell subclusters revealed a significant reduction in the Beta_2 population, one of the 3 most abundant β-cell subclusters in WT islets (**Fig. 8b-e**). Differential gene expression analysis showed that genes upregulated in Beta_2 cells were enriched for pathways involved in protein synthesis and ribosome biogenesis (**Fig. 8f**), whereas downregulated genes were enriched for functions related to membrane potential regulation, ion transport, and synaptic organization (**Fig. 8g**).

**Figure 9.**
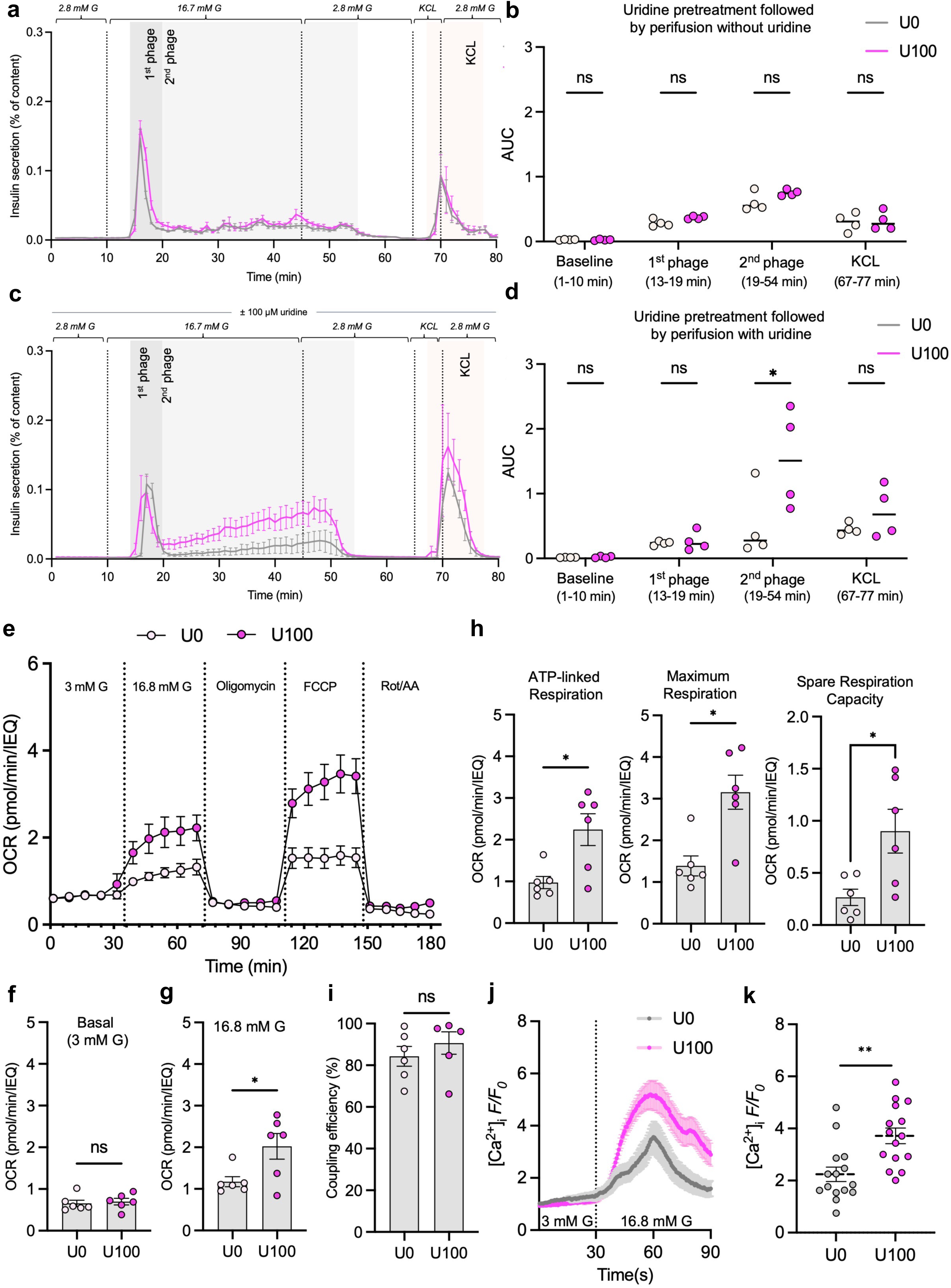
Uridine enhances second-phase glucose-stimulated insulin secretion and mitochondrial oxidative capacity. (a) Insulin secretion from perifused islets with or without uridine pretreatment. Islets in the U100 group were treated overnight with 100 μM uridine, which was omitted throughout the perifusion assay. Insulin secretion was normalised to the total insulin content of 50 islet equivalents (IEQ) and is presented as mean ± SEM. *n* = 4 mouse biological replicates. A consistent 3-min delay in changes in glucose concentration occurred because of the dead volume in the connecting tubing. (b) AUC quantification of the insulin secretory responses shown in (**a**) during the baseline (1-10 min), first phase (13-19 min), second phase (19-54 min), and KCl-stimulated phase (67-77 min). (c) Insulin secretion from perifused islets with or without uridine pretreatment. Islets in the U100 group were treated overnight with 100 μM uridine, which was also included throughout the perifusion assay. Insulin secretion was normalised to the total insulin content of 50 IEQ and is present as mean ± SEM. *n* = 4 mouse biological replicates. (**d**) AUC quantification of the insulin secretory responses shown in (**c**) during the baseline (1-10 min), first phase (13-19 min), second phase (19-54 min), and KCl-stimulated phase (67-77 min). (**e**) Real-time Seahorse analysis of oxygen consumption rate (OCR) in mouse islets cultured overnight with or without 100 μM uridine. OCR was measured under basal conditions and following sequential exposure to glucose and mitochondrial inhibitors and was normalised to islet IEQ. *n = 6* mouse biological replicates. (**f,g**) Mean OCR under basal– and stimulatory-glucose conditions, respectively, calculated from the experiment shown in (**e**). (**h,i**) Mitochondrial functional parameters–including ATP-linked respiration, maximal respiration, spare respiration capacity, and coupling efficiency–calculated from the data shown in (**e**) according to the manufacturer’s definitions. (**j,k**) Ca^2+^ responses in islets transduced with adenovirus-GCaMP5, treated overnight with or without uridine, and exposed sequentially to 3 and 16.8 mM glucose. Representative traces and mean *F/F_0_* between 30 and 90 s from one experiment are shown; *n* = 15 cells per group. Data are presented as mean ± SEM. Statistical significance in (**b**,**d**) was determined by two-way ANOVA followed by Šídák’s multiple-comparison test. Data in (**f-i**) were analyzed using paired two-tailed Student’s *t* tests, and data in (**j,k**) were analyzed using a two-tailed Welch’s *t* test. \**p* < 0.05, \*\**p* < 0.01; ns, not significant.

Pseudo-bulk analysis of β-cells further revealed significant downregulation of multiple genes critical for β-cell function in *Upp1* KO islets, including *Ins1* and *Ins2* (**Fig. 8h**). Genes associated with β-cell identity and maturity exhibited mixed changes, with decreased expression of *Pdx1* and *MafA* but increased expression of *Neurod1* and *Pax6* (**Fig. 8h**). Genes involved in stimulus-secretion coupling were also differentially regulated. Expression of *Slc2a2*, encoding GLUT2, the principal glucose transporter in mouse β-cells, was decreased, whereas expression of *Gck*, encoding glucokinase, the enzyme that initiates glycolysis, was increased (**Fig. 8h**). In addition, *Kcnj11* and *Abcc8*, which encode the subunits of the ATP-sensitive potassium (K_ATP) channel, were differentially expressed in *Upp1* KO β-cells (**Fig. 8h**).

In WT islets, uridine treatment induced coordinated upregulation of genes encoding glycolytic enzymes, components of calcium signaling and the exocytotic machinery, and ribosomal proteins, consistent with the enhanced GSIS following uridine treatment (**Fig. 8i**). Concordant with increased activity in glycolytic activity and insulin granule secretion, genes involved in insulin biosynthesis were also upregulated (**Fig. 8i**). In contrast, these transcriptomic responses were largely abolished or markedly attenuated in uridine-treated *Upp1* KO islets (**Fig. 8i**). Notably, uridine treatment increased, rather than decreased, *Upp1* expression in WT islets (**Fig. 8i**), suggesting that UPP1-mediated uridine metabolism remains active during uridine exposure.

Consistent with uridine-mediated GSIS potentiation, genes encoding components of oxidative phosphorylation, the TCA cycle, pyruvate oxidation, and mitochondrial metabolite transport were upregulated by uridine in WT islets, whereas these responses were abolished in *Upp1* KO islets (**Fig. 8j**). Collectively, these findings indicate that uridine activates transcriptional programs supporting glycolysis, stimulus-secretion coupling, insulin biosynthesis, and mitochondrial oxidative metabolism in a UPP1-dependent manner. These data provide mechanistic context and hypothesis-generating insight but do not fully explain the acute secretory mechanism.

### Uridine enhances second-phase glucose-stimulated insulin secretion and mitochondrial oxidative capacity

To determine how uridine-mediated potentiation affects the dynamics of glucose-stimulated insulin secretion, we performed islets perifusion assays. In the first experiment, islets were pretreated with uridine overnight, but uridine was omitted throughout the perifusion assay, including the 90-min equilibration period. As expected [21], untreated WT islets exhibited a biphasic secretory response to glucose, consisting of a prominent first phase over a fixed period (7 min in our study), followed by a sustained but low and flat second phase that persist throughout glucose stimulation [39] (**Fig. 9a**). Uridine-pretreated islets showed no difference from untreated islets in the AUCs for baseline, first-phase, second-phase, or KCl-stimulated insulin secretion (**Fig. 9b**). This result was consistent with the static GSIS findings in the uridine-withdrawal group (**Fig. 1d**), indicating that the continued presence of uridine is required to maintain uridine-mediated GSIS potentiation.

In a second perifusion experiment, uridine was included throughout the assay for islets that had received uridine during overnight culture. Under these conditions, uridine-treated islets exhibited greater second-phase insulin secretion than control groups, whereas baseline, first phase glucose-stimulated, and KCl-stimulated insulin secretions remained comparable between groups (**Fig. 9c,d**). Notably, reducing the glucose concentration to 2.8 mM after the stimulatory phase rapidly suppressed insulin secretion in both groups (**Fig. 9c**). Thus, uridine amplifies glucose-stimulated insulin secretion without overriding its glucose dependence.

To determine whether uridine-mediated potentiation of GSIS was associated with enhanced glucose metabolism and mitochondrial function, we measured islet oxygen consumption using a Seahorse assay (**Fig. 9e**). Uridine-treated islets exhibited a greater OCR AUC across the complete assay than untreated islets (**ESM Fig. 9c**), whereas basal OCR remained comparable between groups (**Fig. 9f**). The glucose-induced increase in OCR was greater in uridine-treated islets (**Fig. 9g**), indicating an enhanced mitochondrial response to glucose. Uridine also increased ATP-linked respiration and maximal respiration without altering coupling efficiency (**Fig. 9h,i**) or proton leak (**ESM Fig. 9d**). These findings indicate that uridine enhances glucose-responsive oxidative phosphorylation and mitochondrial respiratory capacity without increasing basal respiration, promoting mitochondrial uncoupling, or altering the efficiency with which respiration is coupled to ATP production. Spare respiratory capacity was also greater in uridine-treated islets (**Fig. 9h**), further supporting an increase in mitochondrial reserve capacity. Collectively, these results identify mitochondrial oxidative metabolism as one component of glucose signaling enhanced by uridine.

Thus, uridine amplifies the mitochondrial and metabolic response to stimulatory glucose, which may contribute to the enhanced sustained second phase of GSIS.

Although the AUC for first-phase insulin secretion was not significantly increased by uridine, uridine treated islets consistently exhibited an earlier onset of secretion (**Fig. 9c**). We therefore measured glucose-induced intracellular Ca^2+^ response in islets transduced with adenovirus-GCaMP5. Uridine treated islets exhibited a greater Ca^2+^ response to stimulatory glucose (**Fig. 9j,k**), suggesting that enhanced Ca^2+^ signaling may contribute to the early amplification of GSIS by uridine.

### *In vivo* effects of β-cell specific *Upp1* deletion on glucose tolerance and *ex vivo* effects of chronic uridine exposure and hyperglycemia on uridine-mediated GSIS potentiation

To assess the *in vivo* significance of uridine-UPP1 signal, we performed oral glucose tolerance tests in mice after ablation of β-cell UPP1. Chow-fed *Upp1* KO^β^ mice exhibited impaired glucose tolerance with basal blood glucose concentrations unchanged (**Fig. 10a**). HFD-fed *Upp1* KO^β^ mice showed exacerbated glucose intolerance with elevated basal blood glucose (**Fig. 10c**). Under both dietary conditions, glucose-induced increase in blood insulin was trended to be lower in *Upp1* KO^β^ mice (**Fig. 10b,d**), suggesting potential compensatory change in β-cell function to impaired glucose tolerance by β-cell *Upp1* deletion.

**Figure 10.**
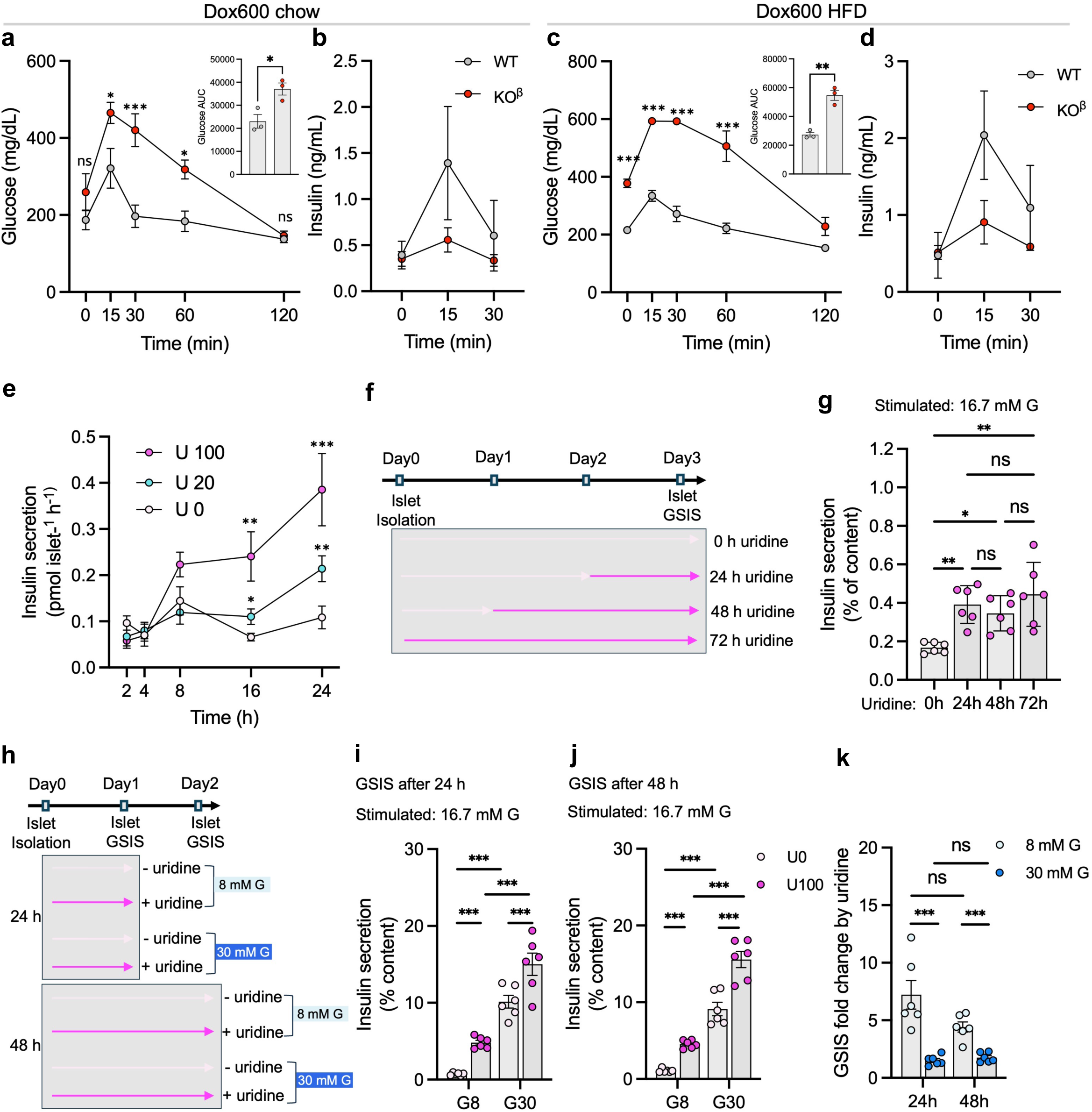
*In vivo* effects of β-cell-specific *Upp1* deletion on glucose tolerance and *ex vivo* effects of prolonged uridine exposure and hyperglycemia on uridine-mediated GSIS potentiation (**a,b**) Blood glucose and plasma insulin concentrations during oral glucose tolerance tests (OGTTs) in control and *Upp1* KO^β^ mice after 3 weeks of Dox-chow feeding. *n* = 3 mice per group. (**c,d**) Blood glucose and plasma insulin concentrations during OGTTs in control and *Upp1* KO^β^ mice after 5 weeks of Dox-HFD feeding. *n* = 3 mice per group. (**e**) Islets from WT mice were cultured in RPMI 1640 containing 8 mM glucose and 0, 20, or 100 µM uridine. Culture medium was collected at the indicated time points. *n* = 5 mouse biological replicates per group. (**f**) Experimental design for assessing the effects of prolonged uridine exposure on GSIS. Freshly isolated WT islets were cultured for 72 h in RPMI 1640 containing 8 mM glucose, with 100 µM uridine present for the indicated durations. The culture medium was replaced daily throughout the culture period. (**g**) Static GSIS in islets subjected to the 72 h culture protocol shown in (**f**). *n* = 6 mouse biological replicates per group. (**h**) Experimental design for assessing the effects of glucose concentration on uridine-mediated GSIS potentiation. Freshly isolated WT islets were cultured for 24 or 48 h in RPMI 1640 containing either 8 or 30 mM glucose, with or without 100 μM uridine. (**i,j**) Static GSIS in islets following 24 and 48 h treatment, respectively, according to the protocol shown in (**h**). *n* = 6 mouse biological replicates per group. (**k**) Uridine-induced fold change in GSIS calculated from the data shown in (**i**,**j**). *n* = 6 mouse biological replicates per group. Data are presented as mean ± SEM. Statistical significance in (**a-d**) was determined by two-way ANOVA followed by Šídák’s multiple-comparison test, whereas glucose AUC in (**a,c**) were analyzed using two-tailed unpaired Student’s t-test. Statistical significance in (**e**) was determined by two-way ANOVA followed by Tukey’s multiple comparisons test; significance in (**g**) was determined by one-way ANOVA followed by Tukey’s multiple comparisons test; and significance in (**i**-**k**) was determined by two-way ANOVA followed by Šídák’s multiple comparisons test. \**p* < 0.05, \*\**p* < 0.01, \*\*\**p* < 0.001; ns, not significant.

UPP1 catalyzes uridine catabolism, raising the possibility that it prevents excessive intracellular uridine accumulation, which might otherwise impair β-cell secretory function. Because the GSIS assays assessed insulin secretion during only 1-h of glucose stimulation following a 1-h preincubation at low glucose, the effect of uridine on longer-term cumulative insulin secretion remained unclear. To address this question, we cultured islets overnight in RPMI 1640 containing 8 mM glucose and 0, 20, or 100 μM uridine. Islets were then transferred to fresh medium containing the corresponding uridine concentrations, and culture medium was collected after 2, 4, 8, 16, and 24 h to measure cumulative insulin secretion. In untreated islets, cumulative insulin secretion began to plateau within 8 h (**Fig. 10e**). In contrast, insulin secretion continued to increase through 24 h in uridine-treated islets, with 100 μM uridine eliciting a greater response than 20 μM uridine (**Fig. 10e**). Islet insulin content remained comparable among the three groups at the end of the 24-h culture period (**ESM Fig. 10a**). Thus, these findings indicate that uridine enhances cumulative β-cell insulin secretion in a concentration-dependent manner under these experimental conditions without depleting islet insulin stores.

To determine the effect of prolonged uridine exposure on GSIS, islets were cultured for 72 h, with 100 μM uridine added during the final 24, 48, or 72 h of the culture period (**Fig. 10f**). Uridine-mediated potentiation of GSIS was comparable across all exposure durations (**Fig. 10g**), and total islet insulin content remained unchanged (**ESM Fig. 10b**). These findings indicate that prolonged exposure to elevated uridine concentrations does not diminish uridine-mediated potentiation of GSIS.

Chronic hyperglycemia, which commonly develops in the setting of insulin resistance, can progressively impair β-cell function through glucotoxicity. To determine whether hyperglycemia affects uridine-mediated potentiation of insulin secretion, islets were cultured for 24 h or 48 h in medium containing either 8 or 30 mM glucose before GSIS analysis (**Fig. 10h**). After 24 h of culture, insulin secretion was higher in uridine-treated islets than in the corresponding untreated controls under both glucose concentrations (**Fig. 10i**). Islet insulin contents was substantially lower after culture in 30 mM glucose than after culture in 8 mM glucose (**ESM Fig. 10c**), suggesting that high-glucose exposure reduced intracellular insulin stores. However, uridine did not alter insulin content under either glucose condition, indicating that uridine potentiated insulin secretion without further depleting β-cell insulin stores.

After 48 h of culture, uridine-mediated potentiation remained detectable under both glucose conditions (**Fig. 10j**). Insulin content in islets cultured in 8 mM glucose declined to a level comparable to that in islets cultured in 30 mM glucose (**ESM Fig. 10d**), suggesting that prolonged culture itself adversely affected insulin production and/or storage. Uridine did not reduce total islet insulin content under either glucose condition (**ESM Fig. 10d**), further indicating that its potentiation of secretion did not occur at the expense of insulin stores. At both 24 and 48 h, uridine responsiveness—calculated as the uridine-induced fold increase in insulin secretion at 16.7 mM glucose—was significantly lower in islets cultured in 30 mM glucose than in those cultured in 8 mM glucose (**Fig. 10k**). However, untreated islets cultured in 30 mM glucose already exhibited substantially elevated GSIS (**Fig. 10i,j**). Thus, the apparent reduction in uridine responsiveness after high-glucose exposure may be partially attributable to the elevated insulin secretion in the absence of uridine, which increases the denominator used to calculate the fold response and is a recognized feature of early adaptive response to glucose stress [40]. In uridine-treated islets, *Upp1* expression was modestly reduced after 24 h culture in 30 mM glucose than in 8 mM glucose but was comparable between two glucose conditions after 48 h (**ESM Fig. 10e**). In the absence of uridine, however, *Upp1* expression was unaffected by 30 mM glucose at either time point (**ESM Fig. 10e**), indicating that the increased GSIS induced by high-glucose culture alone was not associated with altered *Upp1* expression. Collectively, these findings indicate that prolonged exposure to high glucose attenuates, but does not abolish, uridine-mediated potentiation of GSIS.

## Discussion

Pyrimidine metabolism is essential for maintaining the specialized function of pancreatic β-cells, which require a continuous supply of pyrimidine nucleotides to support RNA synthesis, protein translation, phospholipid biosynthesis, membrane biogenesis, and insulin granule production. Despite these fundamental requirements, the role of pyrimidine metabolism in regulating β-cell function under physiological conditions and during diabetes progression remains largely unexplored. To our knowledge, this study provides the first evidence that extracellular uridine potentiates GSIS in a UPP1-dependent manner, identifying β-cell UPP1 as a previously unrecognized regulator that links pyrimidine metabolism to glucose homeostasis.

UPP1 catalyzes the phosphorolytic cleavage of uridine into uracil and ribose-1-phosphate, the latter of which can subsequently enter central carbon metabolism. This mechanism has been proposed to underlie the protective effects of uridine against ischemic injury in cardiomyocytes and neurons [41, 42] and to support the survival of glucose-deprived cancer cells by providing an alternative carbon source [33, 34]. Our findings suggest that β-cells use uridine in a distinct, glucose-dependent manner. Whereas cancer cells preferentially catabolize uridine under glucose-limiting conditions and suppress uridine utilization when glucose is abundant [33, 34], uridine-mediated potentiation of GSIS requires stimulatory glucose concentrations. Uridine alone was insufficient to trigger insulin secretion under low-glucose concentrations; however, in the presence of stimulatory glucose, uridine enhanced second-phase insulin secretion and increased mitochondrial respiratory capacity. These findings indicate that UPP1-mediated uridine metabolism primarily amplifies glucose-stimulated signaling rather than serving as a glucose-independent alternative fuel source. Consistent with this model, UPP1-deficient islets exhibit normal GSIS under conventional uridine-free assay conditions, suggesting that endogenous glucose metabolism remains intact while the uridine-dependent amplification pathway is selectively disrupted. Single-nucleus multiomic analysis further supports this model by demonstrating that uridine enhances glycolytic activity and glucose uptake in β-cells, effects that are abolished or markedly attenuated in UPP1-deficient islets. Interestingly, elevated UPP1 expression has also been reported to promote glycolytic flux in gastric cancer (GC) cells for proliferation [43], suggesting that UPP1-mediated regulation of glycolytic flux may represent a conserved mechanism across diverse cell types despite distinct physiological outcomes.

Our study further identifies the MEK/ERK pathway as a critical downstream mediator of uridine action in glucose-stimulated β-cells. MAPK signaling has long been implicated in nutrient sensing, β-cell adaptation, and insulin secretion [37], although its precise contribution to GSIS remains controversial. Previous studies have shown that inhibition of ERK1/2 has little effect on GSIS in INS-1 cells [38] but partially suppresses GSIS in MIN6 cells and primary rat islets [44], suggesting that the requirement for MAPK signaling is context dependent. Under our experimental conditions, pharmacological inhibition of MEK completely abolished uridine-mediated GSIS potentiation without impairing the islet response to stimulatory glucose alone, indicating that UPP1-dependent activation of the MEK/ERK pathway specifically mediates the amplifying effect of uridine rather than the core glucose-sensing response. However, constitutive MEK activation failed to restore uridine responsiveness in *Upp1* KO islets, indicating that MEK/ERK signaling is necessary but not sufficient for uridine-mediated GSIS potentiation. Thus, additional UPP1-dependent mechanisms are likely required to produce the complete secretory response.

Beyond its acute effects on insulin secretion, ERK signaling contributes to the unique biology of β-cells by mediating glucose-regulated insulin gene transcription [37]. Consistent with this role, UPP1-deficient islets exhibited impaired uridine-induced ERK activation together with reduced expression of *Ins1* and *Ins2*, suggesting that UPP1-mediated ERK1/2 activation contributes not only to insulin exocytosis but also to the maintenance of insulin biosynthetic capacity. These observations further suggest that UPP1 coordinates both the acute and chronic functions of β-cell by coupling pyrimidine metabolism to transcriptional regulation. Based on these findings, we propose that enhanced glucose-responsive mitochondrial metabolism and MEK/ERK signaling jointly contribute to the insulinotropic effect of uridine. However, the molecular link between UPP1-dependent uridine metabolism and ERK activation remains unsolved and represents an important direction for future investigation.

An intriguing finding was the differential preservation of uridine-mediated GSIS potentiation across models of obesity. Uridine responsiveness was maintained in diet-induced obese mice but completely lost in *ob/ob* mice, coinciding with impaired glucose-stimulated ERK activation in *ob/ob* islets. Basal MEK/ERK phosphorylation is reportedly preserved in diet-induced obesity but attenuated in islets from leptin receptor-deficient *db/db* mice [45]. Together, these observations raise the possibility that intact leptin signaling supports glucose– and uridine-induced activation of the MAPK pathway and, consequently, uridine-mediated potentiation of insulin secretion. However, because *ob/ob* mice exhibit more severe metabolic dysfunction than the HFD-fed mice examined here, the respective contributions of leptin deficiency and disease severity cannot be distinguished. Further studies using inducible leptin-deficiency models across defined stages of metabolic deterioration will be required to resolve these possibilities. It also remains unclear whether HFD further alters UPP1 abundance in the context of leptin deficiency. Collectively, our findings place UPP1-mediated pyrimidine metabolism within a broader signaling network integrating nutrient availability, hormonal signaling, and β-cell functional adaptation.

The translational relevance of these findings is supported by studies in human islets. UPP1 protein abundance was positively correlated with donor HbA_1c_, suggesting that UPP1 expression increases during diabetes progression, potentially as a compensatory response to the increased insulin demand. In contrast, the magnitude of uridine-mediated potentiation of GSIS was inversely correlated with donor HbA_1c_, indicating progressive desensitization of diabetic islets to uridine despite elevated UPP1 expression. UPP1 subcellular localization exhibited substantial heterogeneity among individual islets. Altered subcellular localization may influence UPP1 function, and studies involving larger sample sizes and a broader range of diabetes stages will be required to determine the significance of UPP1 subcellular localization in β-cell function and disease progression.

Circulating uridine concentrations are elevated in both obesity [3, 46] and T2D [7, 8], and UPP1 is a key regulator of systemic uridine homeostasis. Together, these findings raise the possibility that impaired systemic UPP1 activity contributes to dysregulated uridine homeostasis, and consequently, to insulin resistance and β-cell dysfunction during diabetes progression. However, the tissues primarily responsible for UPP1-mediated clearance of circulating uridine remain undefined, the temporal dynamics of circulating uridine during diabetes progression are incompletely understood, and the effects of systemic insulin resistance and deteriorating glycemic control on β-cell uridine responsiveness, independently of changes in UPP1, remains largely unexplored. Our findings demonstrate that high-glucose exposure is sufficient to attenuate, but not immediately abolish, β-cell responsiveness to uridine. Thus, the diminished uridine responsiveness observed during diabetes progression may arise, at least in part, as a consequence of hyperglycemia-induced β-cell dysfunction rather than solely representing a primary defect that initiates glycemic deterioration. Conversely, β-cell-specific *Upp1* deletion produced a more evident impairment in glucose tolerance under metabolic stress than under basal conditions, despite no significant alteration in circulating insulin response to glucose. These findings suggest that β-cell UPP1 may be particularly important for supporting adaptive insulin secretion during metabolic stress, although compensatory β-cell responses and systemic insulin clearance may obscure changes in circulating insulin. Thus, dysregulation of UPP1-uridine signaling and β-cell dysfunction may develop as interconnected and mutually reinforcing consequences of chronic hyperglycemia and metabolic stress. Future studies are needed to determine how β-cell UPP1 activity and systemic uridine availability are coordinately regulated during diabetes development and progression. Such studies will help clarify the contribution of impaired UPP1-uridine signaling to islet β-cell dysfunction and worsening glycemic control.

Several limitations should be acknowledged. First, 100 µM uridine was selected for the *in vitro* islet study because it reproducibly potentiated GSIS in WT islets without detectable adverse effects on islet viability and represents one of the lowest concentrations reported for mammalian cell studies ^[16]^. Nevertheless, local uridine concentration within the islet microenvironment may differ substantially from circulating levels because macrophages have been proposed to release uridine ^27^. Direct measurement of interstitial uridine concentrations under physiological and diabetic conditions will therefore be important for defining the biologically relevant range of uridine exposure *in vivo*. Second, although our data demonstrate that UPP1 is required for uridine potentiation of GSIS, the downstream metabolic mechanisms remain unresolved. It will be important to determine whether the responsible metabolites arise from uridine catabolism through ribose-1-phosphate production or from uridine salvage into UTP and CTP, thereby influencing glycolysis, phospholipid synthesis, membrane biogenesis, or nucleotide homeostasis. Finally, because UPP1 is expressed in multiple tissues, it remains unclear how β-cell UPP1 activity is coordinated with systemic uridine metabolism. Identifying the regulators of UPP1 expression and activity, as well as the mechanisms responsible for elevated plasma uridine in obesity and diabetes, will provide a more comprehensive understanding of how pyrimidine metabolism contributes to β-cell adaptation and the pathogenesis of T2D.

## Supporting information

ESM

## Acknowledgements

We thank Qiangrong Liang (New York Institute of Technology) for providing the CaMEK1 adenovirus and Sangeeta Dhawan (City of Hope) for discussion regarding the use of frozen human islets for research. We are also grateful to the Transgenic Technology Center Core at UT Southwestern Medical Center for rederiving the *Upp1* KO mouse line and generating *Upp1*^f/f^ mice. We thank the NIDDK-funded Integrated Islet Distribution Program (IIDP) (RRID:SCR_014387) at the City of Hope, NIH Grant # U24DK098085, and the Breakthrough T1D (formerly JDRF)-funded IIDP Islet Award Initiative for providing human islets for research for this study. Schematic figures were created in BioRender.

## Data and Resource Availability

All data related to this work can be found in the paper or ESM materials.

## Author contributions

Y.P. conceived of and conducted experiments with help from L.D. and Y.W.. Q.X. conducted single-nucleus multiomic analysis. L.F. and C.J. conducted studies involving mouse models with help from I.N-S., K.W., P.E.S., and Z.V.W.; G.L., E.M.M., E.O., L.T., I. J., P. W., A.G-O., and D.T. helped with islet functional studies and human tissue assessments; Y.D. conceived of and designed the study, conducted experiments, and wrote the manuscript. All the authors commented on and approved the manuscript. Y.D. is the guarantor of this work.

