## Supplementary material for "Role of Uridine Phosphorylase 1 (UPP1) in Pancreatic β-Cell Physiology and Its Dysregulation in Obesity and Type 2 Diabetes": ESM

**ESM Table 1.** Characteristics of human islet donors included in the GSIS study.

| Donor | Gender | Age (years) | BMI | HbA1c | Race | Donor type |
|---|---|---|---|---|---|---|
| Hu 1265 | Male | 20 | 38.3 | 37 mmol/mol<br>5.50% | Hispanic | Non-diabetic |
| Hu 1267 | Male | 63 | 32.1 | 43 mmol/mol<br>6.10% | Caucasian | Pre-diabetic |
| Hu 1269 | Female | 36 | 30.3 | 29 mmol/mol<br>4.80% | African American | Non-diabetic |
| Hu 1270 | Male | 45 | 22.8 | 36 mmol/mol<br>5.40% | Hispanic | Non-diabetic |
| Hu 1271 | Male | 34 | 33 | 40 mmol/mol<br>5.80% | Hispanic | Pre-diabetic |
| Hu 1280 | Female | 55 | 31.7 | 41 mmol/mol<br>5.90% | Hispanic | Pre-diabetic |
| Hu 1281 | Male | 61 | 33.3 | 44 mmol/mol<br>6.20% | African American | Pre-diabetic |
| Hu 1284 | Female | 54 | 39.7 | 41 mmol/mol<br>5.90% | Asian | Pre-diabetic |
| Hu 1291 | Male | 20 | 42.7 | 40 mmol/mol<br>5.80% | Hispanic | Pre-diabetic |
| Hu 1292 | Male | 32 | 41.7 | 34 mmol/mol<br>5.30% | Hispanic | Non-diabetic |
| Hu 1293 | Male | 43 | 29.9 | 50 mmol/mol<br>6.70% | Caucasian | Diabetic |
| Hu 1294 | Female | 42 | 33.3 | 37 mmol/mol<br>5.50% | Hispanic | Non-diabetic |
| Hu 1296 | Female | 39 | 36.5 | 42 mmol/mol<br>6.00% | Hispanic | Pre-diabetic |
| Hu 1297 | Male | 56 | 27.7 | 69 mmol/mol<br>8.50% | Hispanic | Diabetic |
| Hu 1300 | Female | 49 | 37.5 | 38 mmol/mol<br>5.60% | Caucasian | Non-diabetic |
| Hu 1301 | Male | 60 | 27.8 | 38 mmol/mol<br>5.60% | Caucasian | Non-diabetic |
| Hu 1314 | Male | 63 | 30.7 | 50 mmol/mol<br>6.70% | Caucasian | Diabetic |
| Hu 1315 | Female | 37 | 26.4 | 33 mmol/mol<br>5.20% | Caucasian | Non-diabetic |
| Hu 1316 | Male | 47 | 29.7 | 55 mmol/mol<br>7.20% | Hispanic | Diabetic |
| Hu 1321 | Female | 53 | 33.3 | 40 mmol/mol<br>5.80% | Hispanic | Pre-diabetic |
| Hu 1322 | Male | 21 | 31.2 | 41 mmol/mol<br>5.90% | Caucasian | Pre-diabetic |
| Hu 1323 | Male | 42 | 26.4 | 37 mmol/mol<br>5.50% | Caucasian | Non-diabetic |
| Hu 1324 | Female | 39 | 31.4 | 32 mmol/mol<br>5.10% | Hispanic | Non-diabetic |

Non-diabetic donors were defined as having an HbA1c ≤ 5.6%; pre-diabetic doners as having an HbA1c of 5.7-6.4%; diabetic doners as having an HbA1c ≥ 6.5%, according to established diagnostic criteria for non-pregnant individuals. The islets were obtained by direct pickup from the Southern California Islet Cell Resource Center (SC-ICRC) at City of Hope.

**ESM Table 2.** Characteristics of human islet donors included in the UPP1 Western blot analysis.

| Donor | Gender | Age | BMI | HbA1c | Donor type | Islet type |
|---|---|---|---|---|---|---|
| SAMN41032378 | M | 36 | 19.3 | 36 mmol/mol<br>5.4% | Non-diabetic | frozen |
| SAMN43474066 | M | 54 | 22.3 | 34 mmol/mol<br>5.3% | Non-diabetic | frozen |
| SAMN50929223 | M | 42 | 23.4 | 34 mmol/mol<br>5.3% | Non-diabetic | frozen |
| SAMN51186627 | M | 45 | 22.7 | 36 mmol/mol<br>5.4% | Non-diabetic | frozen |
| SAMN55389234 | M | 66 | 24.6 | 22 mmol/mol<br>4.1% | Non-diabetic | frozen* |
| SAMN37973608 | F | 55 | 31.7 | 41 mmol/mol<br>5.9% | Non-diabetic | frozen |
| SAMN48457719 | F | 48 | 19.6 | 29 mmol/mol<br>4.8% | Non-diabetic | frozen |
| SAMN55343530 | F | 53 | 34.3 | 26 mmol/mol<br>4.5% | Non-diabetic | frozen* |
| SAMN45223776 | M | 48 | 25.3 | 39 mmol/mol<br>5.7% | Pre-diabetic | frozen |
| SAMN46746834 | M | 58 | 39.8 | 45 mmol/mol<br>6.3% | Pre-diabetic | frozen* |
| SAMN48795793 | M | 53 | 17.5 | 48 mmol/mol<br>6.5% | Diabetic | frozen |
| SAMN55315454 | M | 49 | 35.5 | 54 mmol/mol<br>7.1% | Diabetic | frozen* |
| SAMN56641486 | M | 59 | 33.9 | 51 mmol/mol<br>6.8% | Diabetic | frozen* |
| SAMN40995252 | F | 55 | 32.9 | 51 mmol/mol<br>6.8% | Diabetic | frozen |
| SAMN43565566 | F | 55 | 42.2 | 61 mmol/mol<br>7.7% | Diabetic | frozen |
| SAMN57497796 | F | 49 | 43.9 | 50 mmol/mol<br>6.7% | Diabetic | frozen* |

Non-diabetic donors were defined as having an HbA1c ≤ 5.6%; pre-diabetic doners as having an HbA1c of 5.7-6.4%; diabetic doners as having an HbA1c ≥ 6.5%, according to established diagnostic criteria for non-pregnant individuals. Frozen islets from these donors were used for the experiments shown in Fig. 7A-C. *, indicates islet batches that were received fresh and subsequently snap-frozen in our lab upon receipt. Human islets were obtained through the Integrated Islet Distribution Program (IIDP) via shipment or direct pickup from the Southern California Islet Cell Resource Center (SC-ICRC) at City of Hope.

**ESM Table 3.** Characteristics of human pancreas donors included in the UPP1 immunofluorescence study.

| Donor | Gender | Age | BMI | HbA1c | Donor type |
|---|---|---|---|---|---|
| Hu 958 | M | 24 | 25 | 31 mmol/mol<br>5.0% | Non-diabetic |
| Hu 965 | M | 24 | 50 | 38 mmol/mol<br>5.6% | Non-diabetic |
| Hu1149 | M | 18 | 24 | 29 mmol/mol<br>4.8% | Non-diabetic |
| Hu1156 | M | 44 | 30 | 33 mmol/mol<br>5.2% | Non-diabetic |
| Hu1157 | M | 51 | 24 | 28 mmol/mol<br>4.7% | Non-diabetic |
| Hu 954 | F | 33 | 23 | 30 mmol/mol<br>4.9% | Non-diabetic |
| Hu1152 | F | 58 | 37 | 34 mmol/mol<br>5.3% | Non-diabetic |
| Hu1164 | F | 43 | 32 | 38 mmol/mol<br>5.6% | Non-diabetic |
| Hu1168 | F | 30 | 36 | 26 mmol/mol<br>4.5% | Non-diabetic |
| Hu1198 | F | 46 | 24.4 | 37 mmol/mol<br>5.5% | Non-diabetic |
| Hu 988 | M | 62 | 28 | 42 mmol/mol<br>6.0% | Pre-diabetic |
| Hu1006 | M | 47 | 31 | 45 mmol/mol<br>6.3% | Pre-diabetic |
| Hu1008 | M | 63 | 26 | 41 mmol/mol<br>5.9% | Pre-diabetic |
| Hu1012* | M | 46 | 23 | 45 mmol/mol<br>6.3% | Pre-diabetic |
| Hu1021 | M | 46 | 21 | 40 mmol/mol<br>5.8% | Pre-diabetic |
| Hu 992 | F | 40 | 23 | 39 mmol/mol<br>5.7% | Pre-diabetic |
| Hu 996 | F | 43 | 30 | 39 mmol/mol<br>5.7% | Pre-diabetic |
| Hu1009 | F | 53 | 33 | 42 mmol/mol<br>6.0% | Pre-diabetic |
| Hu1041 | F | 48 | 34 | 39 mmol/mol<br>5.7% | Pre-diabetic |
| Hu1043 | F | 51 | 23 | 40 mmol/mol<br>5.8% | Pre-diabetic |
| Hu 587 | M | 44 | 48.4 | 48 mmol/mol<br>6.5% | Diabetic |
| Hu 902 | M | 61 | 42.1 | 86 mmol/mol<br>10.0% | Diabetic |
| Hu 926 | M | 52 | 32.1 | 50 mmol/mol<br>6.7% | Diabetic |
| Hu1005 | M | 62 | 36 | 57 mmol/mol | Diabetic |

| | | | | | |
|---|---|---|---|---|---|
| | | | | 7.4% | |
| Hu1007 | M | 57 | 35 | 90 mmol/mol<br>10.4% | Diabetic |
| Hu1002 | F | 52 | 40 | 57 mmol/mol<br>7.4% | Diabetic |
| Hu1020 | F | 59 | 42 | 109 mmol/mol<br>12.1% | Diabetic |
| Hu1078 | F | 51 | 43 | 81 mmol/mol<br>9.6% | Diabetic |
| Hu1142 | F | 50 | 35 | 85 mmol/mol<br>9.9% | Diabetic |
| Hu1173 | F | 42 | 40 | 80 mmol/mol<br>9.5% | Diabetic |

Non-diabetic donors were defined as having an HbA1c ≤ 5.6%; pre-diabetic doners as having an HbA1c of 5.7-6.4%; diabetic doners as having an HbA1c ≥ 6.5%, according to established diagnostic criteria for non-pregnant individuals. *, indicates a donor whose pancreatic tissue section showed no detectable insulin or UPP1 immunofluorescence signal. Human pancreatic tissue sections were obtained from the AR-DMRI Laboratory & Translational Service Centers at City of Hope.

**ESM Table 4**. Pancreatic cell markers for islet single-nucleus multiomic study

| Cell type | Abb. | Marker genes |
|---|---|---|
| Alpha cells | α cell | Ttr, Gcg, Irx2, Slc7a2, Wnk3, Nxph1 |
| Beta cells | β cell | Ins1, Ins2, G6pc2, Lapp, Spock2 |
| Delta cells | δ cell | Sst, Rbp4, Slc2a3, Nrsn1, Spock3 |
| PP cells | γ cell | Ppy, Vsig1 |
| Fibroblast | Fibro | Col1a1 |
| Endotheliocyte | Endo | Pecam1, Cdh5, Vwf |
| Duct cell | Duct | Krt19 |
| Macrophages | Macro | Cytip |

#### ESM Figure Legend

##### ESM Figure 1. Islet insulin content in static GSIS assays

(**a-d**) Islet insulin content corresponding to the static glucose-stimulated insulin secretion (GSIS) assays shown in Fig. 1a-d. $n$ = 4 mouse biological replicates in (**a**), and $n$ = 6 mouse biological replicates in (**b-d**). (**e**) Correlation between donor BMI and uridine-induced GSIS potentiation. GSIS fold change by uridine was the relative GSIS in uridine-treated islets. Donor characteristics are provided in ESM Table 1. Data are presented as mean ± SEM. Statistical significance was determined by one-way ANOVA followed by Tukey's multiple-comparison test. Correlation analysis in (**e**) was performed by linear regression. ns, not significant.

##### ESM Figure 2. Generation and validation of whole-body *Upp1* knockout (KO) and inducible β-cell-specific *Upp1* knockout (KO$^{β}$) mice

(**a**) Whole-body *Upp1* KO mice were rederived from frozen sperm. Briefly, *Upp1* heterozygous mice were rederived from *Upp1* KO frozen sperm obtained from the NIH-supported Mutant Mouse Resource & Research Centers (MMRRC), originally deposited by Giuseppe Pizzorno. Heterozygous mice were intercrossed to produce WT, heterozygous, and *Upp1* KO offspring. In the KO allele, exon 5 was deleted. According to the reported study [15], a primer pair (UPase-m-RT-PCR-F and UPase-m-RT-PCR-R) was designed to amplify a 249-bp RT-PCR product from WT and heterozygous mice but not from *Upp1* KO mice. (**b**) Inducible β-cell-specific *Upp1* KO mice were generated using canonical Cre-LoxP strategy. Mice harboring a *Upp1* floxed allele were produced using an embryonic stem cell clone obtained from the Knockout Mouse Project (KOMP) at University of California, Davis. *Upp1*$^{f/f}$ mice were crossed with MIP-rtTA and TRE-Cre mice to generate inducible β-cell-specific *Upp1* KO mice(*Upp1*$^{f/f}$; MIP-rtTA; TRE-Cre), in which *Upp1* deletion was induced by doxycycline (Dox)-containing diet. (**b-c**) Schematic diagrams of the WT, floxed, and KO *Upp1* alleles. In the floxed allele, exon 6 (115 bp) is flanked by two loxP sites. Primer pair F1/R2 amplifies a 245-bp fragment from the WT allele and a 409-bp fragment from the floxed allele using genomic DNA. Cre-mediated deletion of exon 6 generates the KO allele, which yields a 272-bp PCR product with primer pair F1/R3m, whereas the floxed allele yields a 1118-bp product. (**d**) Genotyping PCR using primer pairs F1R2 and F1/R3m on genomic DNA isolated from the liver and islets of control (*Upp1*$^{f/f}$; MIPrtTA) and *Upp1* KO$^{β}$ (*Upp1*$^{f/f}$; MIPrtTA; TRE-Cre) mice after 2 weeks of Dox600 chow. Primer pair F1/R2 generated only the 409-bp floxed-allele PCR product in both liver and islet samples, but not the 245-bp KO allele product (Lanes 1–4), confirming the absence of WT allele in both control and KO$^{β}$ mice. Primer pair F1/R3m generated the 272-bp KO allele PCR product exclusively in islets from *Upp1* KO$^{β}$ mice (Lane 8), but not in islets of control mice (Lane 7) or liver samples from either genotype (Lanes 5–6), confirming β-cell-specific recombination of the *Upp1* allele. The concurrent detection of the 1118-bp floxed allele PCR product in KO$^{β}$ islets (Lane 8) indicates incomplete recombination within whole islet preparations, consistent with β-cell-specific deletion driven by the *Ins1* promoter (MIP-rtTA/TRE-Cre) while non-β endocrine and non-endocrine islet cells retain the unrecombined floxed allele. (**e**) Sequences of primers used in this figure and *Upp1* primers for qPCR analysis.

##### ESM Figure 3. Basal MAPK signaling is unaltered by uridine treatment in WT and *Upp1* KO islets

(**a**) Islets from each mouse were divided into two groups: untreated (U0) and uridine-treated (U100). Uridine treatment was initiated during overnight culture immediately after islet isolation. Following overnight incubation in RPMI 1640 medium containing 8mM glucose, islets were harvested, lysed, and analyzed by SDS-PAGE and immunoblotting for phosphorylated ERK1/2 (p-ERK1/2) and total ERK1/2, as well as phosphorylated MEK1/2 (p-MEK1/2) and total MEK1/2. (**b**) Quantification of the p-ERK1/ERK1, p-ERK2/ERK2, p-MEK1/2-to-MEK1/2, together with total ERK1, ERK2, and MEK1/2 from the experiments shown in **a**. The average value for each measurement in untreated WT islets within each independent experiment was normalised to 1. $n$ = 4 mouse biological replicates per group. Data are presented as mean ± SEM. Statistical significance was determined by two-way ANOVA followed by Šídák's multiple-comparison test. ns, not significant.

**ESM Figure 4. Basal MAPK signaling is unaltered by uridine treatment in control (*Upp1*$^{\mathbf{f/f}}$, MIP-rtTA) and *Upp1* KO$^{\beta}$ (*Upp1*$^{\mathbf{f/f}}$, MIP-rtTA, TRE-Cre) islets**

(**a**) Mice were fed Dox600 chow for 7 weeks to induce β-cell-specific deletion of *Upp1* before islet isolation. Islets from each mouse were divided into two groups: untreated (U0) and uridine-treated (U100). Uridine treatment was initiated during overnight culture immediately after islet isolation. Following overnight incubation in RPMI 1640 medium containing 8mM glucose, islets were harvested, lysed, and analyzed by SDS-PAGE and immunoblotting for phosphorylated ERK1/2 (p-ERK1/2) and total ERK1/2, as well as phosphorylated MEK1/2 (p-MEK1/2) and total MEK1/2. (**b**) Quantification of the p-ERK1/ERK1, p-ERK2/ERK2, p-MEK1/2-to-MEK1/2, together with total ERK1, ERK2, and MEK1/2 from the experiments shown in (**a**). The average value for each measurement in untreated control (*Upp1*$^{f/f}$, MIP-rtTA) islets within each independent experiment was normalised to 1. $n$ = 4 biological replicates per group. Data are presented as mean ± SEM. Statistical significance was determined by two-way ANOVA followed by Šídák's multiple-comparison test. ns, not significant.

**ESM Figure 5. Acute inhibition of MEK1/2 by PD0325901 abolishes uridine-potentiated GSIS in WT islets**

For each mouse, isolated islets were divided into two groups: untreated (U0) and uridine-treated (U100). Uridine treatment was initiated during overnight culture immediately after islet isolation and maintained throughout the GSIS assay. On the day of GSIS, 1 μM PD or vehicle was added during the final 2 h of culture in RPMI 1640 medium and maintained throughout the GSIS assay. (**a**) Glucose-stimulated insulin secretion, expressed as mean ± SE and normalised to (**b**) the insulin content of 10 islets. $n$ = 6 biological replicates per group. Data are presented as mean ± SEM. Statistical significance in **a** was determined by two-way ANOVA followed by Sidak's multiple-comparisons test. **b** was analyzed using a two-tailed unpaired Student's t-test. $^{**}p < 0.01$; ns, not significant.

**ESM Figure 6. Validation of anti-UPP1 antibodies for detecting UPP1 in mouse islets**

(**a,b**) Islets isolated from control (MIPrtTA) and inducible β-cell *Upp1* overexpression (OE; MIPrtTA, TRE-Upp1) mice were used to validate anti-UPP1 antibody from LSBio (**a**) and Proteintech (**b**). Mice were fed on Dox600 chow for 1 week before islet isolation. View 1 and View 2 were generated from the same Odyssey scan of each immunoblot, with different image display minimum and maximum settings to optimize visualization of UPP1 bands at different expression levels. View 1 highlights the UPP1 band in islets from the UPP1-overexpressing mouse, whereas View 2 optimizes visualization of the endogenous UPP1 band in the control mouse. *, UPP1 protein band. (**c,d**) Islets isolated from WT and whole-body *Upp1* KO mice (**c**), and from control (*Upp1*$^{f/f}$; MIP-rtTA) and inducible β-cell-specific *Upp1* KO mice (**d**) were used to validate UPP1 detection by the LSBio anti-UPP1 antibody. (**e**) Islets isolated from control (*Upp1*$^{f/f}$; MIP-rtTA) and inducible β-cell-specific *Upp1* KO mice were used to validate UPP1 detection by the Proteintech anti-UPP1 antibody. The *Upp1*-overexpressing (*Upp1* OE) and corresponding control islet lysate analyzed in panels **a**, **b**, and **e** was obtained from the same mice. The β-cell-specific *Upp1* knockout (*Upp1* KO$^{\beta}$) and corresponding control islet lysate analyzed in **d** and **e** was obtained from the same mouse. GAPDH or β-Actin served as the loading control. (**f,g**) Total uridine pool size in islet cells and culture medium. Cellular uridine pool (a.u.) was normalized to 100 islet equivalents (IEQ). $n = 4$ biological replicates for *ob/ob* islets, with each replicate derived from one mouse, and $n = 4$ biological replicates for WT islets, with each replicate comprising islets pooled from 4 WT mice. Data are presented as mean ± SEM. Statistical significance in (**f,g**)was determined using paired two-tailed Student's *t* tests. $^{**}p < 0.01$; ns, not significant.

**ESM Figure 7. UPP1 expression in human pancreatic islets**

(**a,b**) Correlation between islet UPP1 protein abundance and donor BMI for all donors combined (**a**) or after stratification by sex (**b**). Donor characteristics are provided in **ESM Table 2**. (**c-f**) Donor-level pseudobulk expression of *INS*, *UPP1*, *PDX1*, and *MAFA* calculated from a human pancreatic islet single-cell RNA-seq dataset (**GSE221156**). Donor-level expression was calculated using *Seurat::AggregateExpression* by grouping cells according to donor ID (orig.ident) and aggregating expression from the RNA data layer. Pairwise comparisons among disease states were performed using two-sided Wilcoxon rank-sum tests, with *p* values shown above the plots. (**g–h**) Correlations between donor-level pseudobulk gene expression and donor $HbA_{1c}$. Linear regression lines with 95% confidence intervals are shown. The fitted regression equation, Pearson correlation coefficient (R), and *p* value are displayed in each plot. Colors indicate disease state: nondiabetic (ND), blue; prediabetic (PD), green; and T2D, red. *n* indicates the number of donors.

**ESM Figure 8. UPP1 subcellular localization in human pancreatic islets.**

Representative immunofluorescence images of human pancreatic sections co-stained for insulin and UPP1. Donor information is provided in **ESM Table 3**. $n = 5$ donors per group. Scale bar, 50 μm. An asterisk (*) denotes a donor whose pancreatic tissue section showed no detectable immunofluorescence for either insulin or UPP1. Representative fields from male donors Hu1157 (ND), Hu1006 (Pre-D), and Hu902 (T2D), and female donors Hu954 (ND), Hu1041 (Pre-D), and Hu1142 (T2D) are presented in **Fig. 7d**.

**ESM Figure 9. Single-nucleus multiomic analysis of islet β-cells from WT and *Upp1* KO mice**

(**a**) Experimental design for single-nucleus multiomic analysis. Age- and sex-matched WT and *Upp1* KO mice (n = 3 per genotype) were used for islet isolation. Islets from each mouse were divided into paired untreated and uridine-treated groups. For each condition, 50 islets per mouse were handpicked and pooled according to genotype and treatment, generating four samples: WT untreated, WT uridine-treated, *Upp1* KO untreated, and *Upp1* KO uridine-treated. All samples were processed concurrently according to the same experimental schedule for nuclei preparation and single-nucleus multiomic analysis. This experimental design controlled for inter-animal variability between treatment groups and reduced technical variation associated with nuclei preparation and library construction. We acknowledge that pooling produced one multiomic sample per genotype and treatment condition; therefore, the three contributing mice do not constitute three independent sequencing replicates. Similarly, the large numbers of nuclei analyzed should not be interpreted as a correspondingly large number of independent biological replicates. (**b**) Cellular composition of WT and *Upp1* KO islets determined by single-nucleus multiomic analysis. Cell identities were assigned based on the marker genes listed in **ESM** Table 4. (**c**) AUC of the complete OCR profiles shown in Fig. 9e. (**d**) Mitochondrial functional parameter–proton leak–calculated from the data shown in Fig. 9e according to the manufacturer's definitions. Data are presented as mean ± SEM. Statistical significance in (**c,d**) was analyzed using paired two-tailed Student's *t* tests. ns, not significant.

**ESM Figure 10. Islet insulin content in static GSIS assays assessing the effects of uridine concentration, chronic exposure, and hyperglycemia on uridine-mediated GSIS potentiation**

(**a-d**) Islet insulin content corresponding to the static glucose-stimulated insulin secretion (GSIS) assays shown in Fig. 10e, g, i and j, respectively. $n = 5$ or 6 mouse biological replicates. (**e**) *Upp1* mRNA levels were normalised by 18s rRNA. Islets were treated according to the protocol shown in **Fig. 10h**. $n = 4$ mouse biological replicates. Data are presented as mean ± SEM. Statistical significance was determined by one-way ANOVA followed by Tukey's multiple-comparison test. $^*p < 0.05$, $^{**}p < 0.01$, $^{***}p < 0.001$; ns, not significant.

### ESM Figure 1

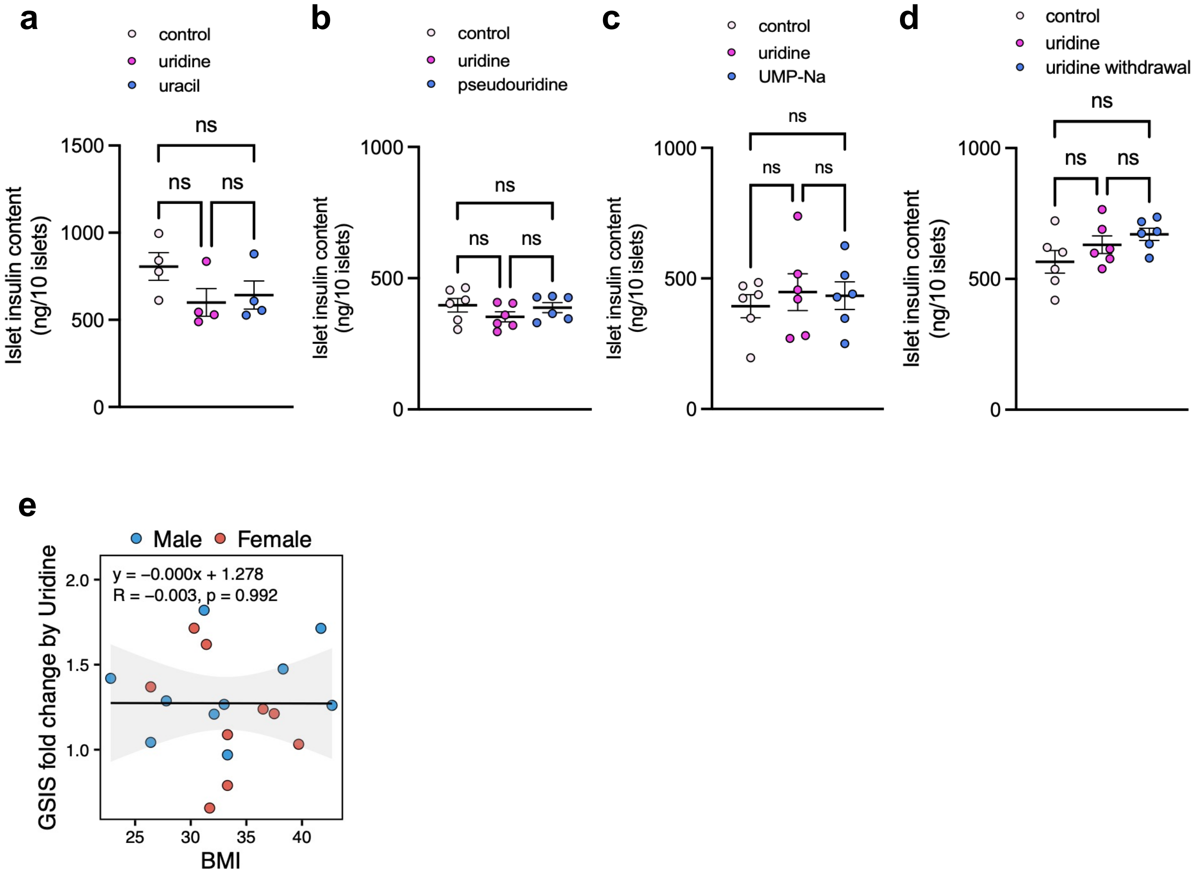
a
control
uridine
uracil
ns
ns
ns
1500
1000
500
0
Islet insulin content
(ng/10 islets)
b
control
uridine
pseudouridine
ns
ns
ns
1000
500
0
Islet insulin content
(ng/10 islets)
c
control
uridine
UMP-Na
ns
ns
ns
1000
500
0
Islet insulin content
(ng/10 islets)
d
control
uridine
uridine withdrawal
ns
ns
ns
1000
500
0
Islet insulin content
(ng/10 islets)
e
Male
Female
y = −0.000x + 1.278
R = −0.003, p = 0.992
2.0
1.5
1.0
25
30
35
40
BMI
GSIS fold change by Uridine

#### ESM Figure 2

**a**

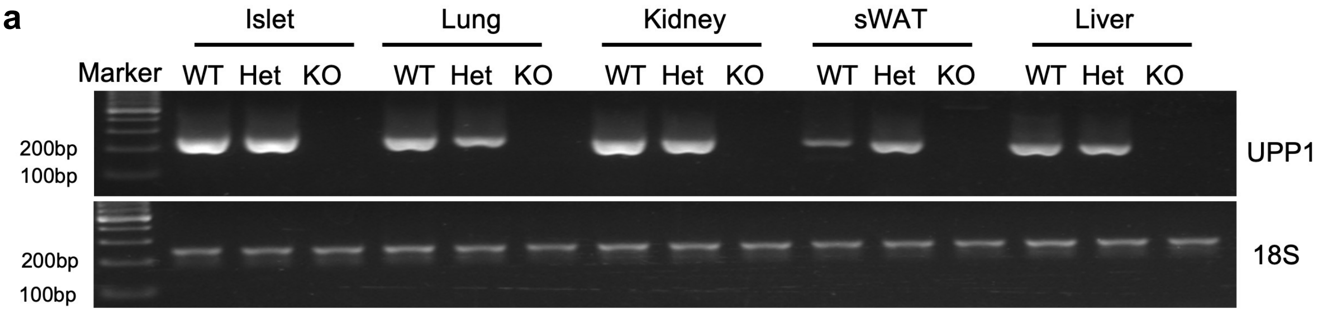


**b**

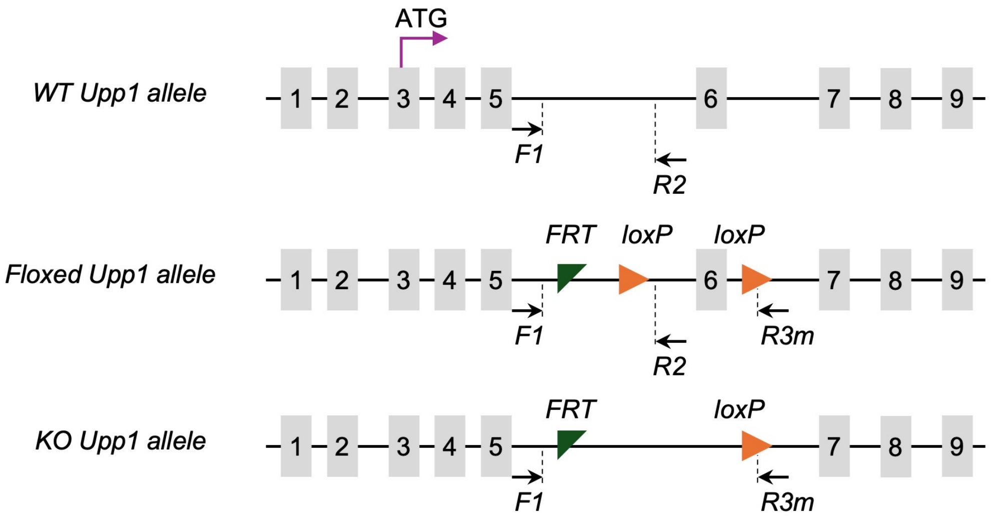


**c**

| Primer set | WT allele | Floxed allele | KO allele |
|---|---|---|---|
| F1/R2 | 245 bp | 409 bp | NA |
| F1/R3m | NA | 1118 bp | 272 bp |

**d**

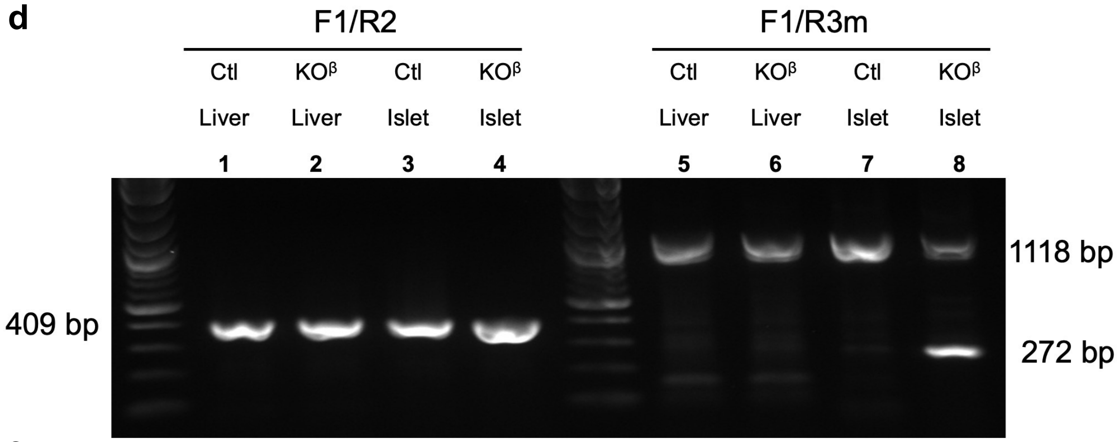


**e**

| Primer pair | Forward (5'-3') | Reverse (5'-3') |
|---|---|---|
| UPase-m-RT-PCR | GCTCTTCCCGGATGAACACC | CGCCTGAAGTGCCAATGC |
| 18S | AGGGTTCGATTCCGGAGAGG | CAACTTTAATATACGCTATTGG |
| F1/R2 | TCAGTGGTAAAACCTTATCCAGTATGGTGGC | CCCACGTGTGGAAGGAGAGTGCTG |
| F1/R3m | TCAGTGGTAAAACCTTATCCAGTATGGTGGC | ATTTGAACTGATGGCGAGCTCAGACC |
| Upp1 qPCR | CCAACATCTGTGCAGGCACT | AACTCCGGCTTGAAGCACTC |

ESM Figure 3

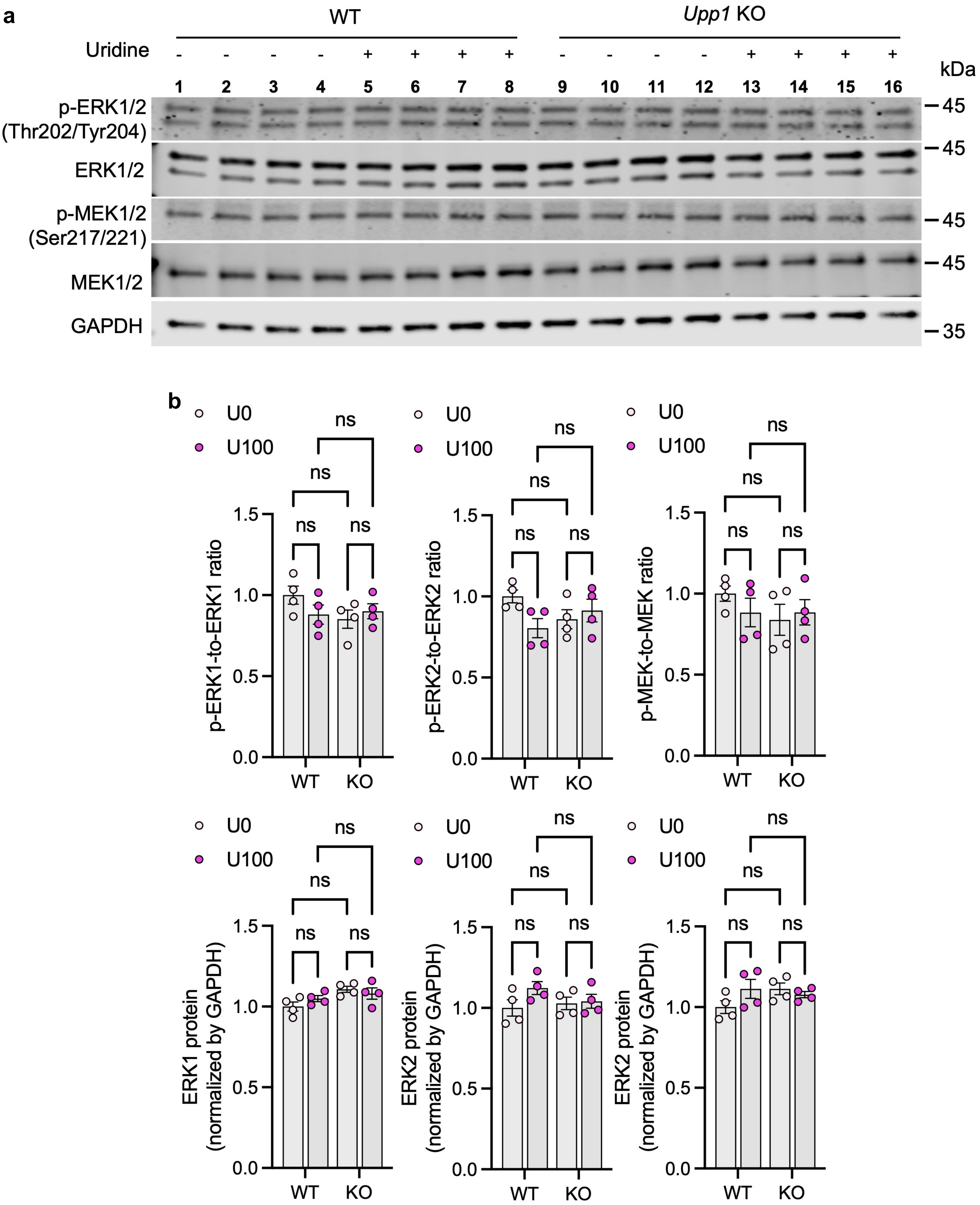

a
WT
Upp1 KO
Uridine
- - - - + + + + - - - - + + + +
1 2 3 4 5 6 7 8 9 10 11 12 13 14 15 16
kDa
p-ERK1/2 (Thr202/Tyr204)
ERK1/2
p-MEK1/2 (Ser217/221)
MEK1/2
GAPDH
45
45
45
45
35
b
U0
U100
ns
p-ERK1-to-ERK1 ratio
p-ERK2-to-ERK2 ratio
p-MEK-to-MEK ratio
ERK1 protein (normalized by GAPDH)
ERK2 protein (normalized by GAPDH)
0.0
0.5
1.0
1.5
WT
KO

ESM Figure 4

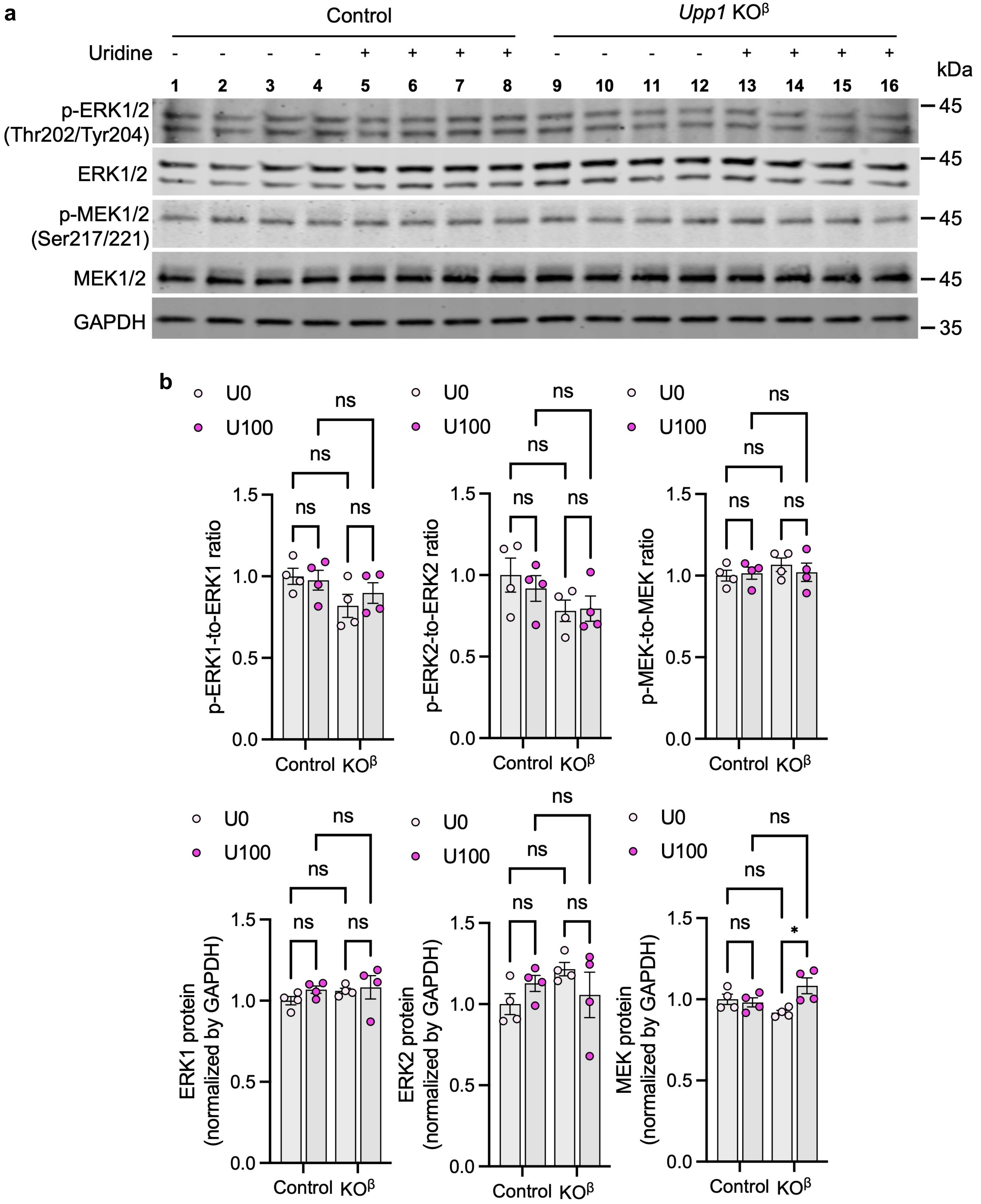
a
Control
Upp1 KOβ
Uridine - - - - + + + + - - - - + + + +
1 2 3 4 5 6 7 8 9 10 11 12 13 14 15 16
kDa
p-ERK1/2 (Thr202/Tyr204)
ERK1/2
p-MEK1/2 (Ser217/221)
MEK1/2
GAPDH
45
45
45
45
35
b
U0
U100
ns
*
p-ERK1-to-ERK1 ratio
p-ERK2-to-ERK2 ratio
p-MEK-to-MEK ratio
ERK1 protein (normalized by GAPDH)
ERK2 protein (normalized by GAPDH)
MEK protein (normalized by GAPDH)
0.0
0.5
1.0
1.5
Control
KOβ

ESM Figure 5

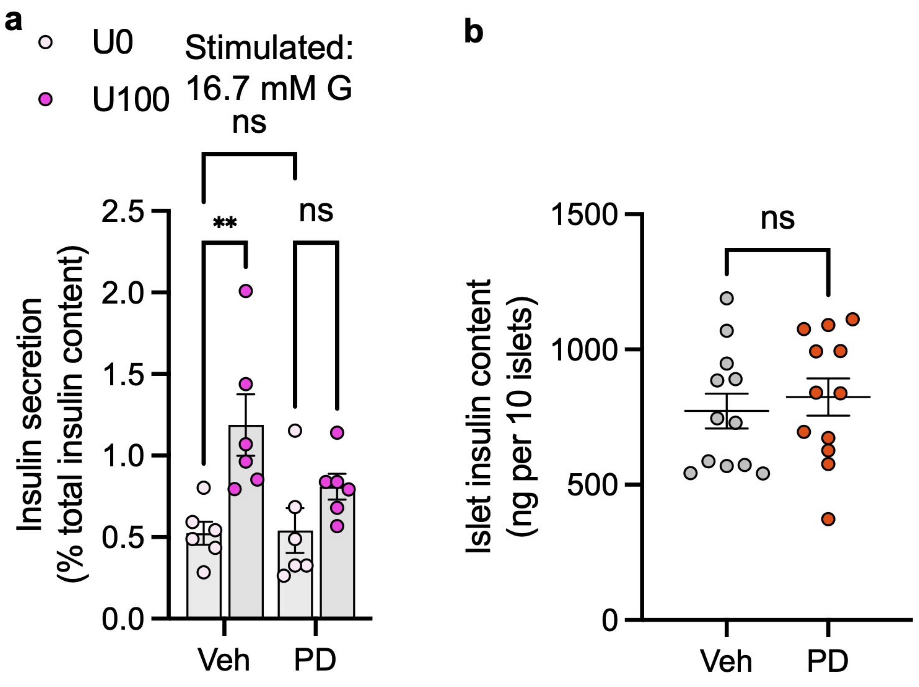
a
U0
U100
Stimulated:
16.7 mM G
ns
**
ns
Insulin secretion
(% total insulin content)
2.5
2.0
1.5
1.0
0.5
0.0
Veh
PD
b
ns
Islet insulin content
(ng per 10 islets)
1500
1000
500
0
Veh
PD

ESM Figure 6

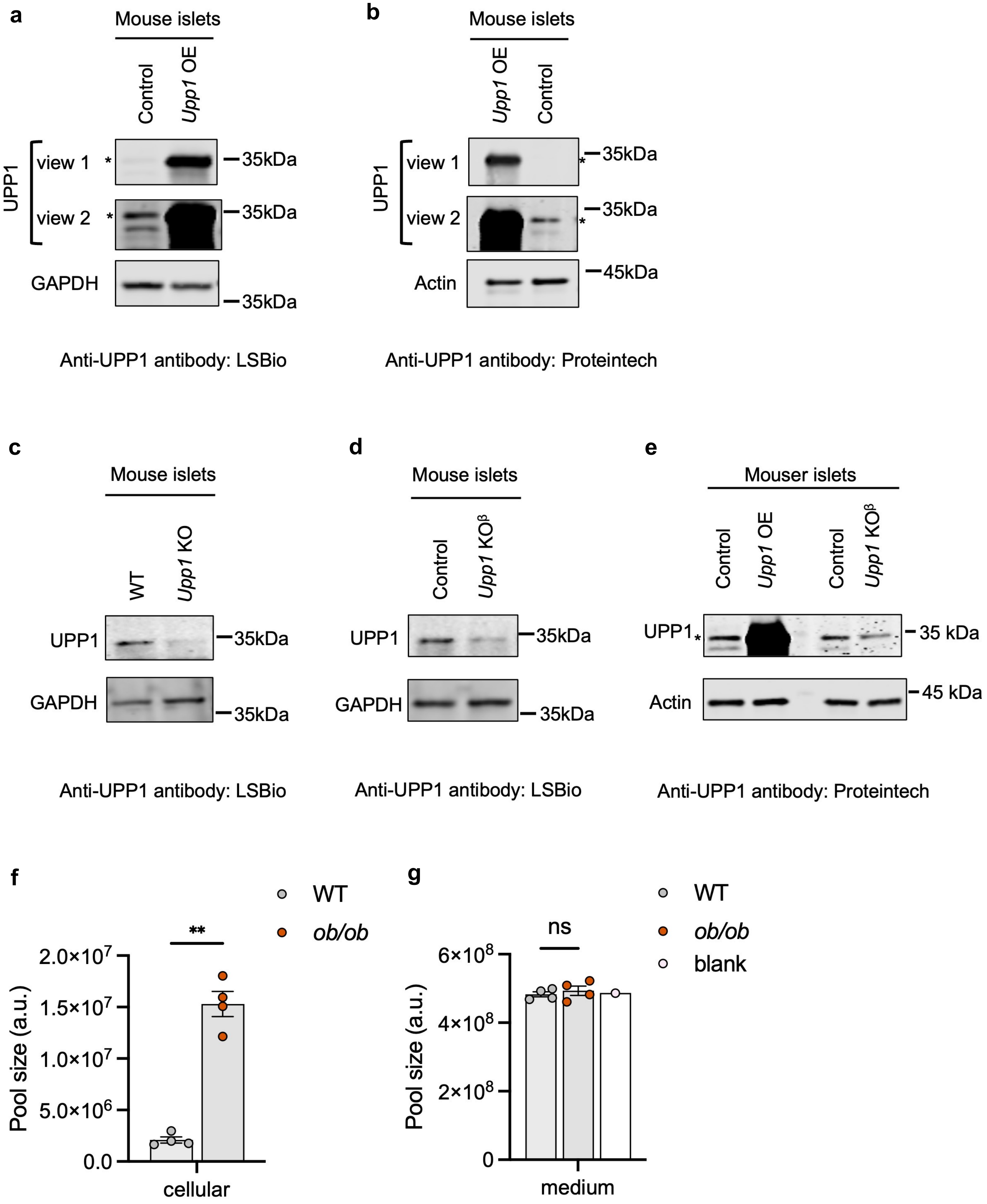
a
Mouse islets
Control
Upp1 OE
UPP1
view 1
view 2
GAPDH
35kDa
35kDa
35kDa
Anti-UPP1 antibody: LSBio
b
Mouse islets
Upp1 OE
Control
UPP1
view 1
view 2
Actin
35kDa
35kDa
45kDa
Anti-UPP1 antibody: Proteintech
c
Mouse islets
WT
Upp1 KO
UPP1
GAPDH
35kDa
35kDa
Anti-UPP1 antibody: LSBio
d
Mouse islets
Control
Upp1 KOβ
UPP1
GAPDH
35kDa
35kDa
Anti-UPP1 antibody: LSBio
e
Mouser islets
Control
Upp1 OE
Control
Upp1 KOβ
UPP1
Actin
35 kDa
45 kDa
Anti-UPP1 antibody: Proteintech
f
WT
ob/ob
**
Pool size (a.u.)
2.0×10^7
1.5×10^7
1.0×10^7
5.0×10^6
0.0
cellular
g
WT
ob/ob
blank
ns
Pool size (a.u.)
6×10^8
4×10^8
2×10^8
0
medium

### ESM Figure 7

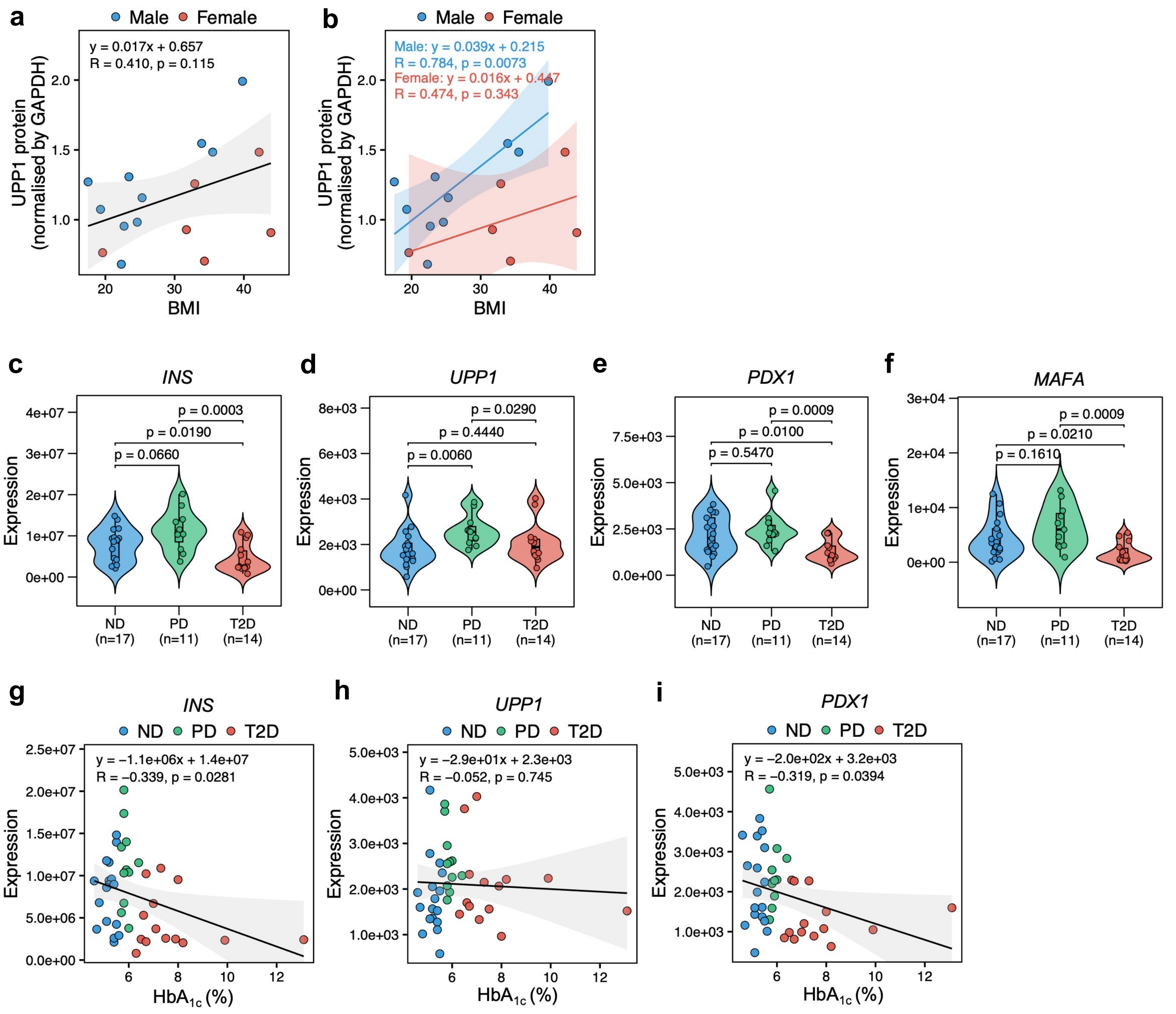
a
Male Female
y = 0.017x + 0.657
R = 0.410, p = 0.115
UPP1 protein (normalised by GAPDH)
BMI
b
Male Female
Male: y = 0.039x + 0.215
R = 0.784, p = 0.0073
Female: y = 0.016x + 0.447
R = 0.474, p = 0.343
UPP1 protein (normalised by GAPDH)
BMI
c
INS
p = 0.0003
p = 0.0190
p = 0.0660
Expression
ND (n=17) PD (n=11) T2D (n=14)
d
UPP1
p = 0.0290
p = 0.4440
p = 0.0060
Expression
ND (n=17) PD (n=11) T2D (n=14)
e
PDX1
p = 0.0009
p = 0.0100
p = 0.5470
Expression
ND (n=17) PD (n=11) T2D (n=14)
f
MAFA
p = 0.0009
p = 0.0210
p = 0.1610
Expression
ND (n=17) PD (n=11) T2D (n=14)
g
INS
ND PD T2D
y = −1.1e+06x + 1.4e+07
R = −0.339, p = 0.0281
Expression
HbA1c (%)
h
UPP1
ND PD T2D
y = −2.9e+01x + 2.3e+03
R = −0.052, p = 0.745
Expression
HbA1c (%)
i
PDX1
ND PD T2D
y = −2.0e+02x + 3.2e+03
R = −0.319, p = 0.0394
Expression
HbA1c (%)

#### ESM Figure 8

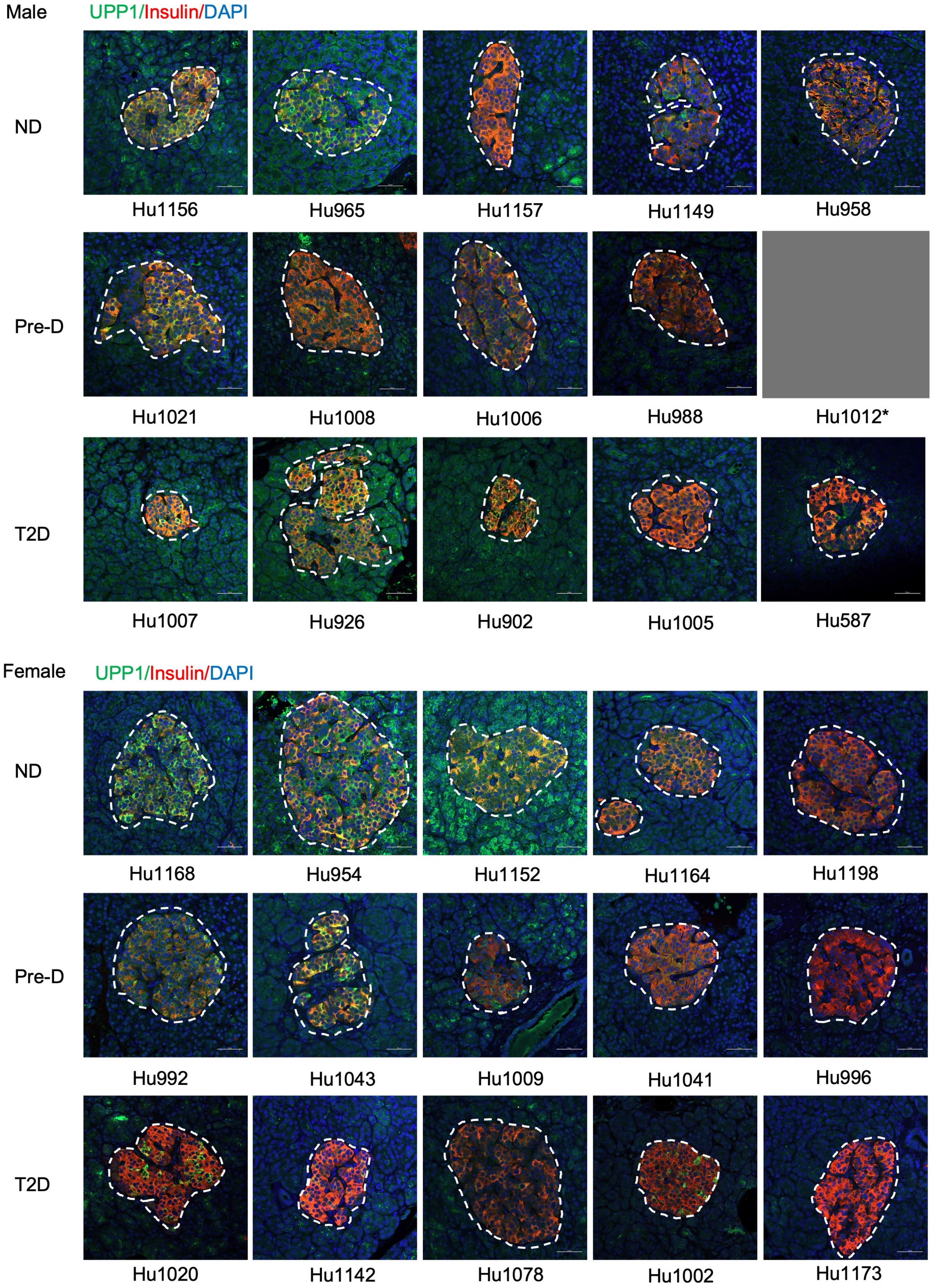
Male
UPP1/Insulin/DAPI
ND
Hu1156
Hu965
Hu1157
Hu1149
Hu958
Pre-D
Hu1021
Hu1008
Hu1006
Hu988
Hu1012*
T2D
Hu1007
Hu926
Hu902
Hu1005
Hu587
Female
UPP1/Insulin/DAPI
ND
Hu1168
Hu954
Hu1152
Hu1164
Hu1198
Pre-D
Hu992
Hu1043
Hu1009
Hu1041
Hu996
T2D
Hu1020
Hu1142
Hu1078
Hu1002
Hu1173

### ESM Figure 9

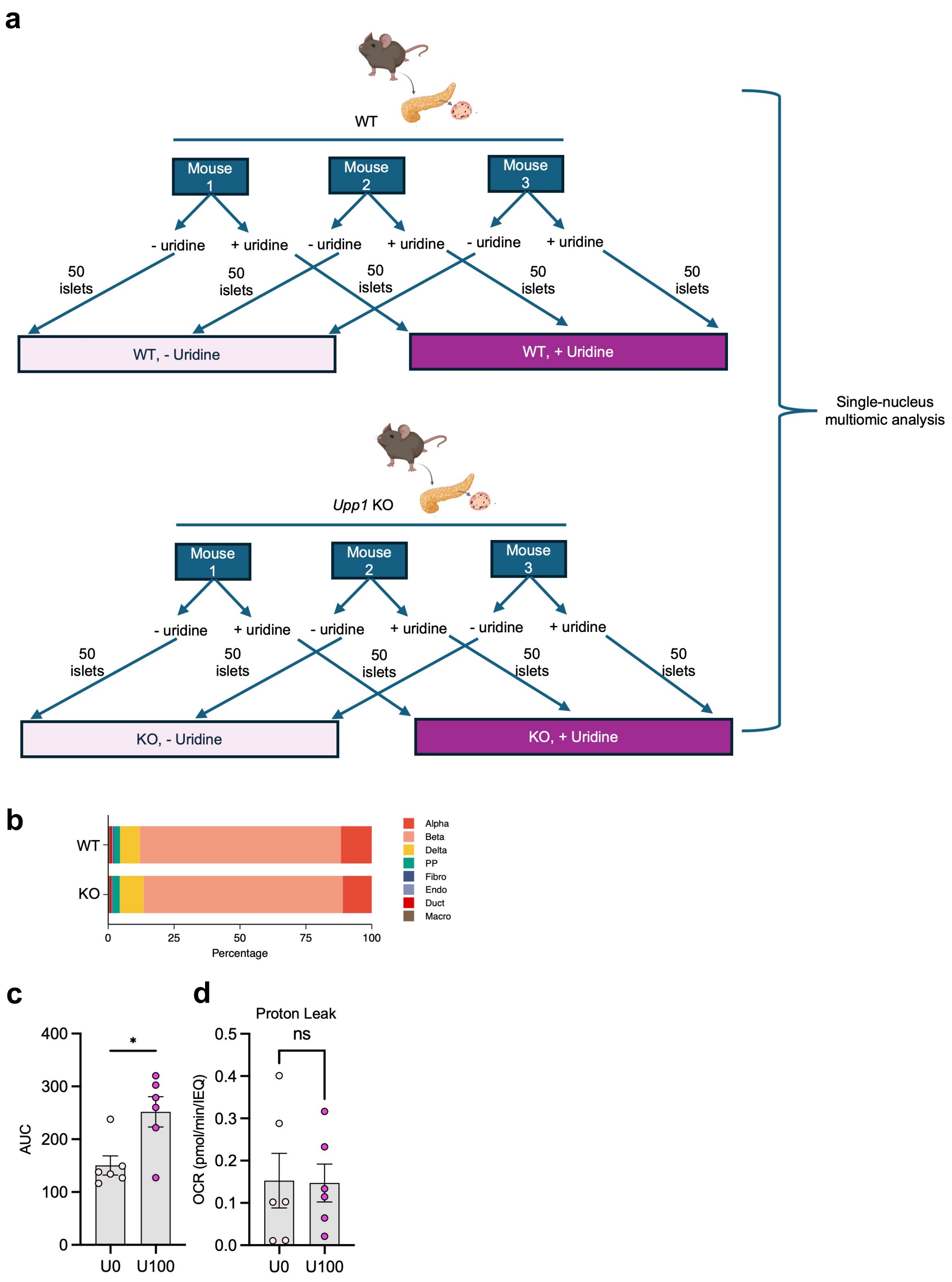
a
WT
Mouse 1
Mouse 2
Mouse 3
- uridine
+ uridine
50 islets
WT, - Uridine
WT, + Uridine
Single-nucleus multiomic analysis
Upp1 KO
KO, - Uridine
KO, + Uridine
b
WT
KO
Percentage
Alpha
Beta
Delta
PP
Fibro
Endo
Duct
Macro
c
AUC
U0
U100
d
Proton Leak
ns
OCR (pmol/min/IEQ)
U0
U100

ESM Figure 10

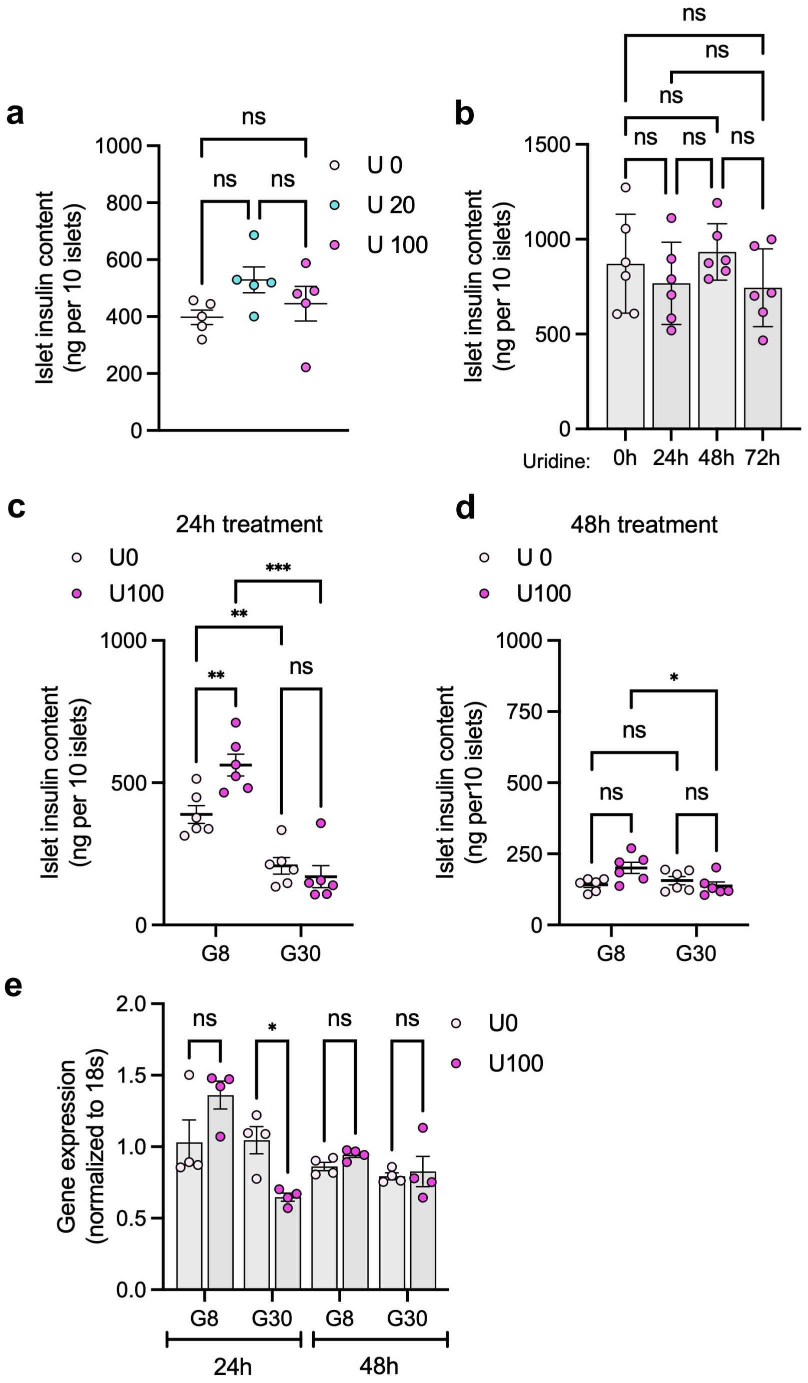

**ESM Method**

**Immunoblotting**

Protein concentrations were determined using the BCA assay (Thermo Scientific). Equal amounts of protein (20 μg per sample) were resolved on 4-20% Tris-glycine gels (Bio-Rad) and transferred to nitrocellulose membranes (Li-Cor). The following primary antibodies were used: GAPDH (Fitzgerald, 10R-G109A), β-actin (Sigma, A5316), UPP1 (LSBio, LS-C374243), UPP1 (Proteintech, 14186-1-AP), phospho-MEK1/2 (S217/S221; Cell Signaling Technology, 9154), MEK1/2 (Cell Signaling Technology, 4694), phospho-ERK1/2 (T202/Y204; Cell Signaling Technology, 9106), ERK (Cell Signaling Technology, 4695). IRDye 800CW goat anti-rabbit (Li-Cor, 925-32211) and IRDye 680RD goat anti-mouse (Li-Cor, 926-68070) secondary antibodies were used for detection.

**Immunofluorescence**

Paraffin sections were deparaffinized, subjected to antigen retrieval (Dako), and blocked with PBS containing 1% BSA and non-immune donkey serum. Sections were incubated with mouse anti-insulin (Thermo Scientific, 14-9769-82) and rabbit anti-UPP1 (Proteintech, 14186-1-AP) primary antibodies, followed by Alexa Fluor 488- or Alexa Fluor 594-conjugated secondary antibodies (Invitrogen). Slides were mounted with ProLong™ Gold Antifade Mountant containing DAPI (Invitrogen, P36931), sealed with glass coverslips, and imaged using a Zeiss LSM 900 confocal microscope with a 20x objective unless otherwise indicated.

**Single-nucleus RNA-seq/ATAC-seq multiomic analysis**

**Sample preparation**

Mouse islets isolated from female WT and Upp1 KO C57BL/6N mice (n = 3 each) were cultured overnight for 16h in complete RPMI 1640 medium with or without 100 μM uridine supplementation. The islets were dispersed into single cells, and all samples were ensured to have >90% viability to minimize dead cell carryover in sequencing. Single-cell suspension samples were processed into nuclei according to the 10x Multiome ATAC + Gene Expression (GEX) protocol (CGOOO338). Samples were collected and washed with PBS (with 0.04% BSA), lysed with pre-chilled lysis buffer for 4 min, washed three times with wash buffer, and resuspended with 10x nuclei buffer at 3,000–5,000 nuclei per μL. Nucleus samples were processed using the Chromium 10x genomics instrument, with a target cell number of 7,000–1,0000 per sample. The 10x Single Cell Multiome ATAC + Gene Expression v1 kit was used according to the manufacturer's instructions for library preparations. A tape station was applied to check the quality of all single-nucleus RNA and ATAC libraries. Sequencing of the RNA and ATAC libraries was performed using the NovaSeq 6000 System (Illumina) and submitted as GSE304559.

**Raw data processing and quality control**

Multiomic sequenced files (fastq files) were processed for demultiplexing and analyzed using *Cell Ranger ARC v2.0*. Genes were mapped and referenced using mouse reference genome *mm10*. Datasets were analyzed using *Seurat 4.4.0* and *Signac 1.11.0* in *R version 4.4.2*. For ATAC data, peaks were called using MACS2 and the genomic positions were mapped and annotated with reference mouse genome *EnsDb.Mmusculus. v79_2.99.0* and *mm10*. Dead cells were removed by excluding cells with very low RNA yields. Low-quality cells including multiplets. Dead cells and poor sequencing depth cells were removed by filtering out cells with low RNA counts (nCount_RNA <3000) and low ATAC counts (nCount_ATAC <2,000); high RNA counts (>25,000) and high ATAC counts (>70,000), and mitochondrial percent > 25.

##### Joint snRNA-seq and snATAC-seq workflow

RNA counts were normalized with SCTransform [1]. Principal component analysis (PCA) was performed on RNA, and UMAP was run on the first 50 principal components (PCs). The ATAC counts were normalized with term-frequency inverse-document-frequency (TFIDF). Dimension reduction was performed with singular value decomposition (SVD) of the normalized ATAC matrix. The ATAC UMAP was created using the second through the $50^{th}$ LSI components.

##### Weighted nearest neighbor analysis

The weighted nearest neighbor (WNN) graph was determined with Seurat's *FindMultiModalNeighbors* to represent a weighted combination of both modalities. The first 50 dimensions of the harmony-corrected RNA reduction and the second through the $50^{th}$ dimensions from the harmony-corrected ATAC reduction were used to create the graph.

##### Annotation of cell types and subclusters

Cell type clusters were identified using weighted snRNA and snATAC modalities. Clusters were identified using WSNN (Weighted Shared Nearest Neighbors) and the SLM algorithm with a resolution = 0.8. Twenty-four clustered were obtained and manually reviewed and annotated into eight cell types according to their canonical markers (**ESM Table 4**). We then used *FindSubCluster* to find subclusters of each cell type. The resolution (0.3, 0.3) was adjusted for each cell type.

##### Differentially expressed genes (DEGs) analysis and gene ontology (GO) enrichment analysis

Differential gene expression analysis was performed using the Seurat package. Genes that were differentially expressed between clusters or specified groups of cells were identified using the *FindAllMarkers* function with the Wilcoxon rank-sum test as the statistical method. For this analysis, we used a 'one-versus-rest' strategy to systematically perform differential gene expression analysis, comparing each cluster to all the other clusters. Only genes expressed in at least 10% of cells in either group and with a log-fold change (logFC) above 0.25 were considered significant. P-values were adjusted for multiple testing using the Bonferroni correction to control the false discovery rate (FDR). The top DEGs were selected based on their adjusted p-values (Padj < 0.05) and ranked by their absolute logFC values. GO enrichment analysis was conducted on the identified DEGs using the *clusterProfiler* R package [2]. Genes from DEGs were used as input for functional enrichment analysis. The analysis included Biological Process (BP), Molecular

Function (MF), and Cellular Component (CC) categories. Enriched GO terms were considered significant if they met the threshold of Padj < 0.05 after multiple testing correction.

##### Average Expression Heatmap Analysis for Pseudo-bulk analysis of β-cells

For heatmap visualization, average gene expression was calculated using the AggregateExpression function in Seurat. Normalized SCT expression values were aggregated across nuclei within each sample group defined by orig.ident, either across all beta cells or within individual beta-cell subclusters. The resulting average expression matrix was scaled by gene and visualized as a heatmap.
